# Pervasive bacterial and prophage hybridization during chronic gut inflammation

**DOI:** 10.64898/2026.05.27.728231

**Authors:** Marta Lourenço, Elsa Seixas, Patrícia Mexia, Minia Antelo-Varela, Patrícia Morais, Karina B. Xavier, Nelson Frazão, Isabel Gordo

## Abstract

Inflammatory bowel diseases (IBD) are modulated by microbiota composition, host genetics, and environmental factors^1^. Humans and other mammals are colonized by multiple strains of *Escherichia coli*, a species that expands in abundance in IBD patients^2^. The state of chronic inflammation characteristic of IBD is expected to intensify selective pressures on the gut microbial community, leading to distinct adaptive trajectories among its constituents. Here we couple *in vivo* experimental evolution with short- and long-read sequencing to test this hypothesis at the level of mutation and horizontal gene transfer (HGT). By colonizing IL10KO mice, a model of IBD^3^, and healthy wild-type mice with two strains of *E. coli*, we show that the *tempo* and *mode* of evolution are strain-specific and strongly shaped by host inflammatory status. Unique mutations associate with host inflammatory status independently of microbiota composition, and rates of transfer are diagnostic of chronic inflammation. Extensive transduction events occur in the inflamed gut, giving rise to hybrid clones that form a new genetic lineage, one that becomes dominant in this disease context. The high levels of recombination between prophages uncovered here point to a critical role of HGT and viral evolution in IBD.

---

Inflammatory bowel diseases (IBD) are complex disorders associated with a myriad of factors, including host genetics, environment, diet, and alterations in the gut microbial community. These diseases are typically characterized by recurrent episodes of intestinal inflammation driven by an atypical immune response^1^. How such fluctuating conditions affect the evolution of gut commensals is poorly known^4^. Proliferation of *E. coli* and reduced microbiota diversity are typically observed in the inflamed gut^5^. However, understanding the adaptive trajectories of *E.coli* evolving under such ecological conditions is still in its infancy^6,7^.

IL10KO mice are widely used to study the pathogenesis of spontaneous, immune-mediated, chronic intestinal inflammation^8,9^. These mice, lack interleukin-10 (IL-10), an anti-inflammatory cytokine produced by monocytes and lymphocytes^10^, important for intestinal immune regulation.

Under conditions of chronic stress, bacteria are exposed to host molecules and metabolites from other microbes. Some can induce SOS responses, which in turn potentiate horizontal gene transfer (HGT)^11^. Host inflammatory responses such as colitis have been shown to alter phage communities in the gut, indicative of higher levels of phage excision^12^.

In this study we use the IL10KO mouse model, in the presence (specific pathogen-free, SPF) and absence (germ-free, GF) of a complex microbiota, to follow the evolutionary dynamics of two phylogenetically distinct *E. coli* strains and to determine how they are affected by chronic inflammation. In these strains evolutionary change occurs by mutational and HGT events, particularly plasmid and phage transfer^13,14^. One strain, MG1655, a human gut isolate that has been adapted to laboratory conditions; belongs to phylogroup A and is hereafter referred to as strain A. The other, a mouse gut isolate capable of donating DNA to strain A, belongs to phylogroup B1, hereafter referred to as strain B1. Using short and long-read sequencing of clones evolving in WT and IL10KO mice, we find that adaptive mutations are host specific and that HGT is a dominant mechanism of evolution in the inflamed gut. The HGT events observed, diagnostic of a stressful gut environment, cause significant changes in the genome and proteome of *E. coli* and result in the emergence of a new lineage capable of reaching high abundances in the inflamed intestine. Furthermore, we find remarkable examples of prophage diversification under chronic inflammation. Overall, our data shows that bacteria and virus diversification is a potential powerful biomarker of IBD.

## Chronic inflammation influences the ecology of *E. coli* strains in the gut

To explore how gut inflammation affects the eco-evolutionary dynamics of gut commensal strains, we used an *in vivo* experimental evolution model of *E. coli* gut colonization of IL10KO and WT mice, with (specific pathogen-free, SPF) or without (germ-free, GF) a complex microbiota. The different host conditions vary in their levels of gut inflammation, as assessed by faecal concentrations of lipocalin-2 (Fig.1a,b; Supplementary Tables 1,2) and IgA (Fig.1c,d, Supplementary Tables 3,4), two common biomarkers of IBD^15–17^. IL10KO SPF mice showed significantly higher levels of lipocalin-2 (LMM, with background and time as fixed factors, P= 0.0015 before colonization and P= 2.74x10^-7^ after colonization, Supplementary Table 2) and IgA (LMM, P = 0.0013 after *E. coli* colonization, Supplementary Table 4) than WT SPF mice, as expected. After colonization with the commensal *E. coli* strains, lipocalin-2 and IgA concentrations were also significantly higher in IL10KO GF than WT GF mice (LMM for mouse genetic background, P=2.64x10^-6^, and P=8.72x10^-4^, Supplementary Tables 1,3, respectively), with lipocalin-2 levels differing by more than two orders of magnitude and IgA levels by more than one order of magnitude. Thus, under both microbiota conditions, IL10KO mice constitute a robust model of chronic inflammation, with microbiota composition having a substantial impact on the inflammatory process.

We next assessed the influence of chronic inflammation on *E. coli* population dynamics (Fig. 1e,f; Supplementary Fig.1). The overall abundance of *E. coli* was higher in IL10KO than in WT SPF mice (LLM with repeated measures, F=33.052, df=1, P=3.3×10⁻⁸), a result that was dependent on the presence of microbiota (LLM with repeated measures, F=0.92, df=1, P=0.34). The two *E. coli* strains stably coexisted in IL10KO mice regardless of microbiota complexity (Supplementary Fig.1). In contrast, coexistence was lost in 2 out of 6 WT SPF mice, in which strain A went extinct after 100 and 300 days post-colonization, respectively (WT1 and WT2; Fig. 1e, f; Supplementary Tables 5,6).

**Figure 1.**
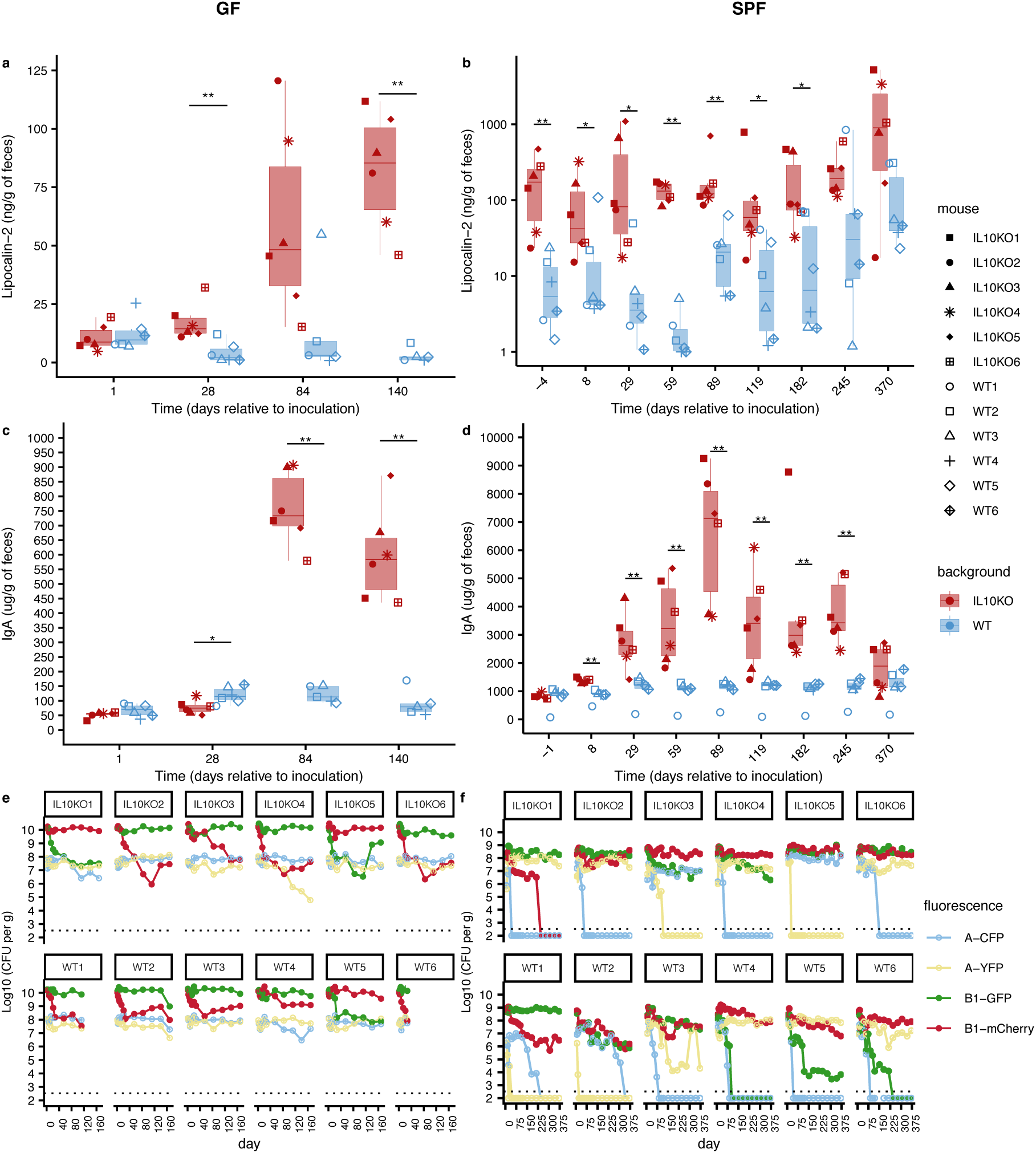
Eco-evolutionary dynamics of *E. coli* in inflamed versus non-inflamed gut. Faecal inflammatory markers and strain dynamics in 6 WT and 6 IL10KO mice colonized with two *E. coli* strains. **a,c,** Boxplots showing faecal lipocalin-2 (LCN2; ng/g faeces) and IgA (µg/g faeces) concentrations measured by ELISA along time in germ-free (GF) WT (blue) and IL10KO (red) mice. **b,d,** Equivalent measurements in specific pathogen-free (SPF) mice along time. More detailed LCN2 data for SPF mice are shown in Table S2. **e, f,** Time series of the relative abundances of the two *E. coli* strains, each carrying one of four fluorescent markers (strain A: CFP and YFP; strain B1: mCherry and sfGFP), in GF (e) and SPF (f) mice. The dotted line indicates the limit of detection for *E. coli*. Note that in WT SPF mice WTM1 and WTM2, strain A went extinct at 100 and 300 days post-colonization, respectively.

While host genetic background influenced the ecological dynamics of the strains (Supplementary Fig.1), the bacterial genomic background had the greatest impact on their evolutionary dynamics. In both IL10KO and WT SPF mice, strain A lost diversity at its neutral fluorescence marker in all mice, whereas strain B1 maintained diversity in 9 out of 12 mice (Fig. 1f). In GF mice, both neutral fluorescence markers were maintained throughout colonization for both strains, and their frequency dynamics consistent with a mode of evolution involving competition and intense clonal interference (Fig. 1e). Thus, in the absence of microbiota, maintenance of neutral diversity was pervasive, whereas the presence of other species in the gut reduced neutral diversity in *E. coli*.

## Higher rate of evolution and more genomic adaptative targets in the inflamed gut

To determine the evolutionary events occurring in each *E. coli* strain in WT and IL10KO mice and to quantify their rates of molecular evolution, we performed pool sequencing of clones from each strain at multiple timepoints throughout colonization. The number of mutational targets was higher in IL10KO than in WT mice irrespective of strain or microbiota complexity (Fig. 2a; Wilcoxon rank-sum test: U=3, P=0.0163 for SPF strain A; U=5, P=0.037 for SPF strain B1; U=0, P=0.0039 for GF strain A; U=2, P=0.01 for GF strain B1). Interestingly, under a complex microbiota, strain B1, despite being a mouse gut isolate, accumulated mutations in a higher fraction of its genomic targets than strain A, including the emergence of mutator subpopulations in both IL10KO and WT background (Fig. 2a, Supplementary Table 8). In the absence of microbiota, the opposite pattern was observed. Mutations accumulated during gut colonization produced a clear separation of the mutational landscapes of each strain according to the host genetic background (Supplementary Fig. 2a), suggesting that the mutational profile of gut commensals may serve as a new biomarker of chronic inflammatory status.

**Figure 2.**
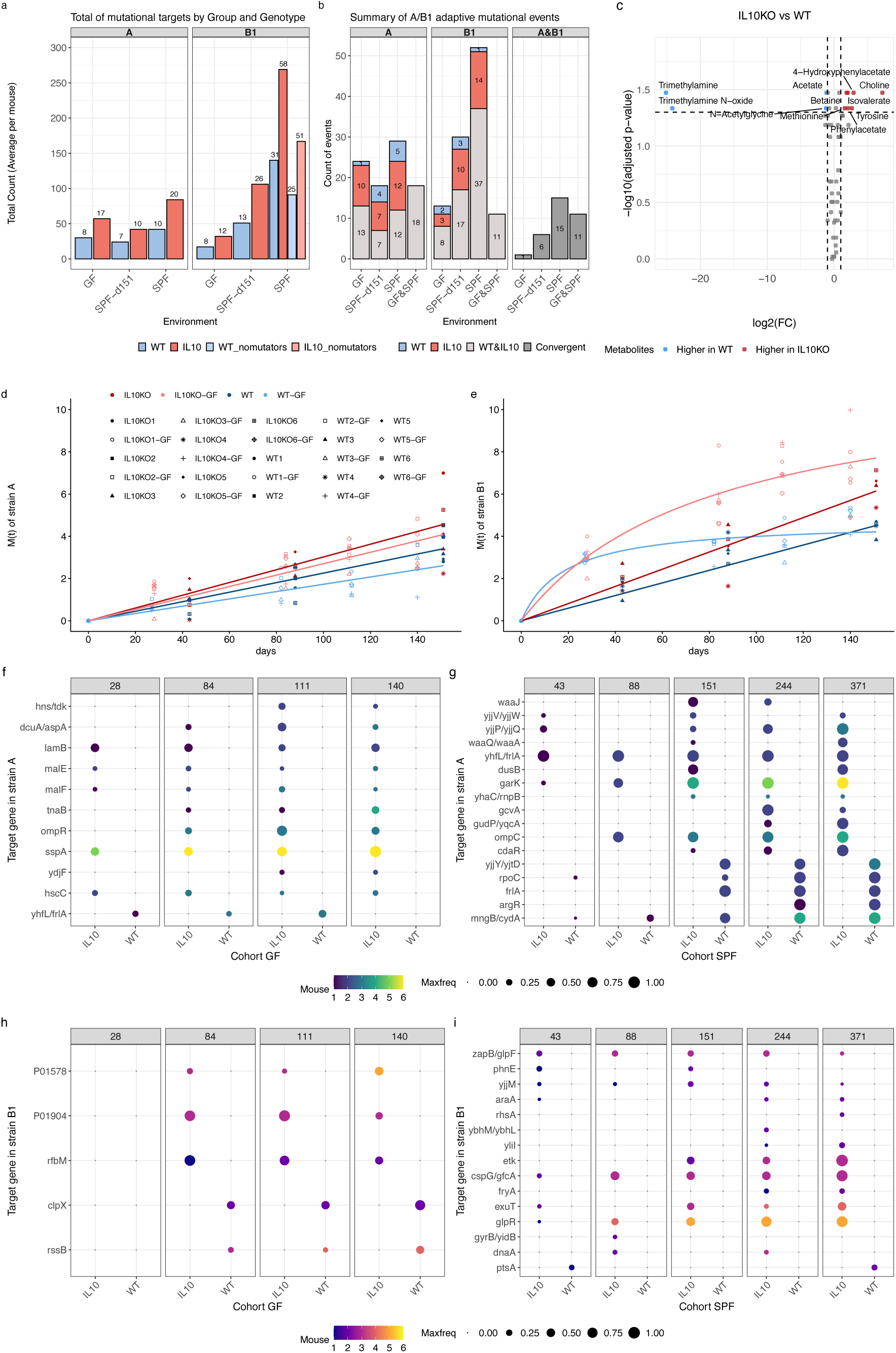
Unique mutational signature of *E. coli* in the inflamed gut. **a,** Total number of mutational targets emerging in *E. coli* strains A and B1 during evolution in WT (blue) and IL10KO (red) mice, shown separately for GF and SPF conditions. Numbers above bars indicate the average number of mutational targets per mouse. Due to time scale differences between GF and SPF mice, data from SPF is shown also for day 151. **b,** Summary of adaptive mutational targets, defined as genes or intergenic regions mutated in more than one mouse, specific to WT (blue), specific to IL10KO (red), shared across both mice cohorts (light grey), or convergent between strains (dark grey). **c** Metabolomic comparison of IL10KO and WT SPF mice, showing metabolites significantly enriched in the inflamed (red) or non-inflamed (blue) gut (p_adj_ = 0.034-0.046). **d, e,** Mutation accumulation over time. Data points represent measured values of M(t), the cumulative sum of allele frequencies at each timepoint. Linear or hyperbolic models were fitted to strain A and strain B1 data, respectively. **f,** Adaptive targets emerging specifically in strain A in GF mice according to host background (healthy versus inflamed). **g,** Adaptive targets emerging specifically in strain A in SPF mice according to host background. **h, i,** Equivalent representations for strain B1 in GF (h) and SPF (i) mice.

We next characterized the genomic targets underlying adaptation to each mouse model (Fig. 2b). We defined *bona fide* adaptive targets as genes or intergenic regions mutated in more than one mouse in which the new allele reached at least 10% frequency among sampled clones. Across all SPF mice, 12 adaptive targets were detected in strain A independent of the host background. Twelve were found specific to the IL10KO cohort and 5 were specific to WT mice (Supplementary Table 7). In strain B1, 37 targets were adaptive across mouse backgrounds. Importantly, we found 14 targets specific to IL10KO and only 1 specific to WT mice (*ptsA*; Supplementary Table 8). Across all GF mice, 13 adaptive targets were detected in strain A. Ten targets specific to IL10KO mice and 1 (*yhfL/frlA* intergenic region) specific to the WT mice were seen (Supplementary Table 9). In strain B1, 8 targets were adaptive across mouse backgrounds. Three were found specific to the IL10KO cohort (*rfbM*, P01578, P01904) and 2 specific to WT mice (*clpX*, *rssB*; Supplementary Table 10). Taken together, these results show that more adaptive mutations arise in IL10KO mice independently of strain identity or microbiota composition, implying that chronic inflammation not only increases the number of emerging mutations but also their convergence across individuals.

To quantify the rate of evolution by mutation in each host, we calculated M(t): the sum of allele frequencies at each sampling timepoint. For strain A, mutation accumulation followed a linear function in both WT and IL10KO backgrounds, in the presence and absence of microbiota, indicating that the rate of mutation accumulation per genome per day (K_A_) remained constant and did not decline over thousands of generations of evolution (Fig. 2d; Supplementary Fig.2b; Supplementary Tables 11,12). Importantly, K_A_ was higher in IL10KO than in WT mice (GF: K_A_=0.027 ± 0.001, P<0.0001; SPF: K_A_=0.030±0.001, P<0.0001 for IL10KO; GF: K_A_=0.017±0.001, P<0.0001; SPF: K_A_=0.024±0.001, P<0.0001 for WT). Thus, for the less dominant strain, mutations accumulated faster in IL10KO than in WT mice over the same period, independently of microbiota.

For the dominant strain B1, mutation accumulation was consistent with a constant rate (K_B_) in the presence of a complex microbiota (K_B_=0.041±0.002, P<0.0001 for IL10KO; K_B_=0.030±0.001, P<0.0001 for WT) but was better described by a hyperbolic function when *E. coli* was the sole colonizer of the gut. This indicates that the evolutionary rate of this commensal strain declines in the absence, but not in the presence, of a complex microbial community (Fig. 2e; Supplementary Fig. 2b; Supplementary Tables 11,12).

## Chronic inflammation leaves a unique mutational signature in *E. coli* genomic adaptation

To understand the influence of chronic inflammation on mutation dynamics, we mapped the frequency trajectories of each mutation in each strain and mouse cohort using *p-τ* plot^18^, which allow different modes of selection to be distinguished (Supplementary Figs. 2c and 3-6; Supplementary Tables 7-10).

**Figure 3.**
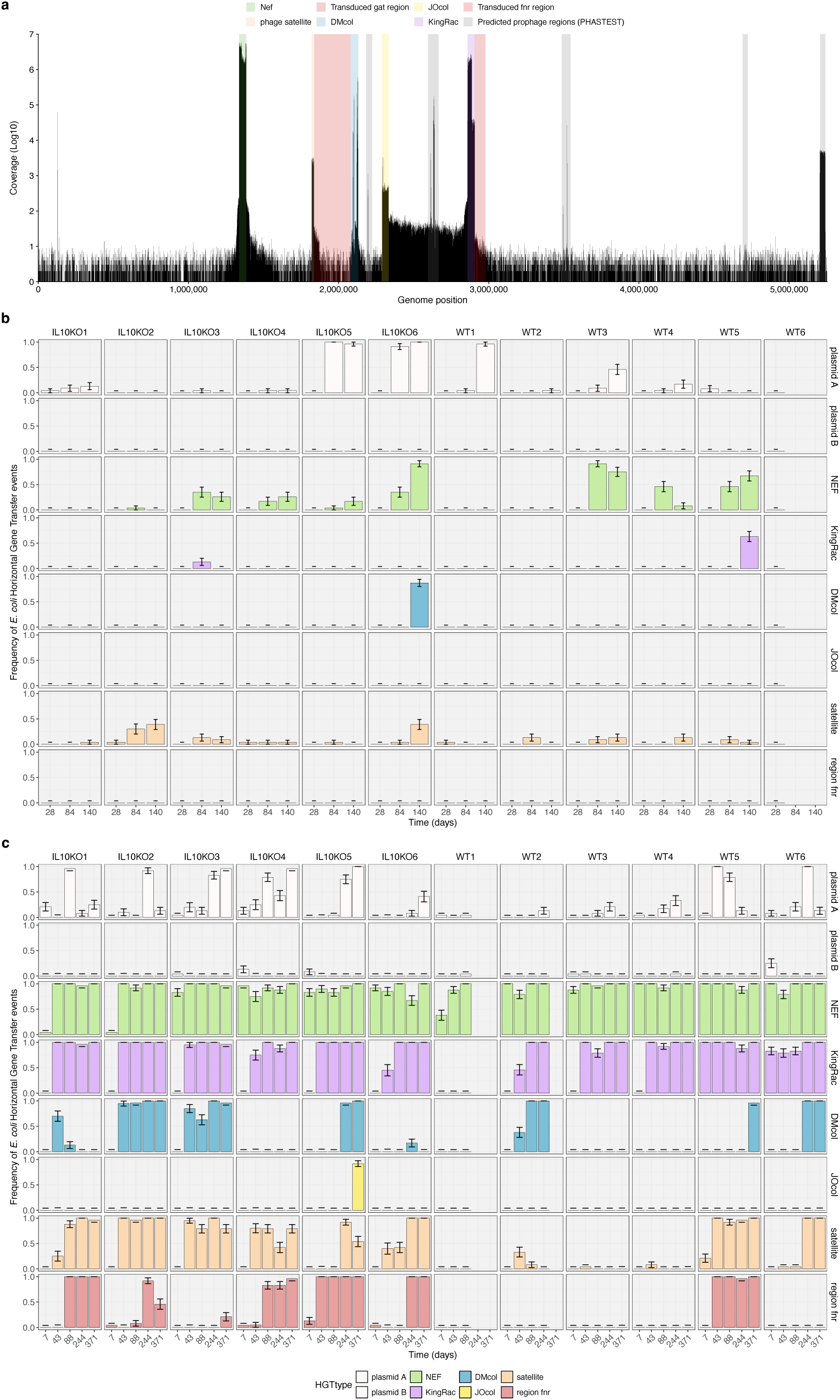
Chronic inflammation in the presence of microbiota heightens HGT events. **a,** Genomic regions of strain B1 with potential for phage-driven transfer, detected by a mitomycin assay *in vitro* followed by sequencing of DNA in the released phage particles (coverage in black). Highlighted in grey are the regions predicted bioinformatically to code for prophages. Prophages or phage satellite effectively transferred from strain B1 to strains A in green, blue, purple, yellow and orange; transduced chromosomal regions in red. **b,c,** Regions transferred *in vivo*, discovered by PCR typing and sequencing of evolved clones from GF (b) and SPF (c) mice.

In strain A, adaptation to GF IL10KO mice was dominated by diversifying selection (Supplementary Fig.2c and 3, Supplementary Table 9), with most mutations maintained at intermediate frequencies. Ten genes were targeted in parallel, among which *sspA*, encoding a stringent starvation response protein, mutated independently and specifically in all IL10KO mice (Fig. 2f; Supplementary Table 9). Remarkably, a single amino acid change (N22S) was responsible for this IL10KO-specific adaptation; while the functional consequence of this substitution is not yet known, serine residues are common phosphorylation targets, suggesting it may alter SspA activity. Notably, SspA interacts with RssB to influence stability of the sigma factor RpoS^19^, a phenotype that is highly polymorphic in *E. coli*^20^. Additional parallel mutations were identified in *ompR*, a DNA-binding transcriptional dual regulator, and in *hscC*, a chaperone known to negatively regulate the activity of sigma factor σ⁷⁰. Mutations specific to IL10KO mice were also found in metabolic pathways, including genes of the maltose metabolism operon (*malF*, *malE*, and *lamB*). In GF WT mice, evolution was similarly dominated by diversifying selection (Supplementary Figs. 2c,3, Supplementary Table 9), though only one parallel adaptive target was detected, in the intergenic region between *yhfL* and *frlA*.

In the presence of a complex microbiota, strain A evolution was dominated by directional selection in both IL10KO and WT mice (Supplementary Fig. 2c,4; Supplementary Table 7). Adaptive targets specific to IL10KO mice included *garK*, encoding a glycerate kinase, which was mutated in all IL10KO mice, as well as *cdaR*, a regulator of glucarate and galactarate metabolism, and *gudP*, a putative D-glucarate and galactarate transporter, each mutated in two or more mice (Fig. 2g; Supplementary Table 7). Interestingly, several mutations potentially affecting the outer membrane were found specifically in the inflamed gut, involving the *waa* operon (important for LPS biosynthesis), *mraY* (important for cell wall peptidoglycan biosynthesis), and *yjjQ* (a transcriptional repressor of genes involved in capsule formation, among other processes). Adaptation specific to WT SPF mice involved 5 targets (Fig. 2g, Supplementary Table 7) previously identified in other experimental evolution studies^13,18,21,22^. Mutation frequency trajectories in strain B1 were consistent with directional selection and clonal interference during evolution in GF IL10KO mice (Supplementary Figs. 2c,5; Supplementary Table 9). Three specific adaptive targets were identified in this cohort: one gene of unknown function and 2 genes involved in colanic acid biosynthesis. In GF WT mice, strain B1 adaptation was consistent with clonal interference, with *clpX* and *rssB* as specific adaptive targets (Fig 2h, Supplementary Table 9), as previously reported^23–25^. As observed for strain A (Supplementary Fig. 7a), several adaptive targets were shared between mouse genetic backgrounds in strain B1 (Supplementary Fig. 7c), including genes (*lacI* and *lacY*) involved in the metabolism raffinose, a highly abundant sugar in GF animals^23,25^, and in the transport of mono-, di-, and tri-saccharides (*ompG*).

In the presence of a complex microbiota and the highly fluctuating inflammatory environment of IL10KO mice, strain B1 mutation frequencies were more consistent with diversifying than directional selection in mice IL10KO2, IL10KO4, and IL10KO6 (Supplementary Figs. 2c, 6bdf). Maintenance of both fluorescent markers at high frequencies in IL10KO2 and IL10KO6 (Fig. 1f) further supports the dominance of diversifying selection in these mice. In mouse IL10KO4, despite large changes in marker frequency (Fig.1f, Supplementary Fig. 6d), a mutator allele emerged late in colonization (Supplementary Table 8) and clonal interference was detected across 5 targets, suggesting intense selective competition that maintains polymorphism at multiple loci. Strain B1 adaptation to SPF IL10KO mice involved mutations in 14 targets; the most ubiquitous (found in 5 out of 6 mice) was *glpR*, a transcriptional repressor of the glycerol-3-phosphate regulon. Intergenic mutations upstream of *glpF*, a glycerol facilitator that permits glycerol diffusion across the inner membrane, further suggest metabolic adaptation to glycerol under inflammatory conditions. Parallel mutations in *exuT*, specific to IL10KO SPF mice, suggest additional metabolic adaptation to hexuronates such as D-galacturonate and D-glucuronate. Consistent with this evolutionary signature, comparison of the metabolomes of IL10KO and WT SPF mice revealed significant differences (Fig. 2c, Supplementary Fig. 8, Supplementary Table 13). In the inflamed gut, choline, 4-hydroxyphenylacetate, betaine, isovalerate, tyrosine, phenylacetate, and methionine were significantly more abundant than in the non-inflamed gut (p_adj_ = 0.034-0.046), a pattern consistent with increased proteolytic fermentation and altered amino acid metabolism driven by intestinal mucosal damage^26,27^. In WT mice, acetate, trimethylamine (TMA), trimethylamine N-oxide (TMAO), and N-acetylglycine were more abundant (p_adj_ = 0.034-0.046; Fig. 2c).

Beyond metabolism, mutations specific to IL10KO SPF mice suggest adaptation *via* modifications of the cell envelope, including mutations in inner membrane proteins GfcA and Etk, which belong to the same operon implicated in group 4 capsule production in some *E. coli* strains^28^ (Fig. 2i). Overall, the specific mutational landscape of adaptation to the inflamed gut points to alterations in both metabolism and membrane composition in strains A and B1.

## Global adaptive targets in *E. coli* independent of the level of gut inflammation

Mutational analysis revealed adaptive targets to the gut environment in each strain independently of inflammatory status or microbiota composition. Strain A evolved a nutritional preference for fructoselysine *via* mutations in 3 genes of the corresponding metabolic pathway: *frlR* (repressor), *frlA* (transporter), and *frlC* (epimerase). These mutations occurred across different mouse genetic backgrounds and microbiota compositions^13,24^ and are driven by the host diet^29^.

Two large deletions (of 87,305 bp and 85,944 bp) were landmarks of global gut adaptation in strain B1, both flanked by an intact tRNA and a partial tRNA, suggesting a high probability of recombination-mediated origin^14^. Other adaptive targets were pervasive across both WT and IL10KO contexts, though their parallelism depended on microbiota composition (Supplementary Figs. 7c, d). Genes *malT* and *chiX* were under selection in strain A when strain B1 was its sole competitor, whereas *melR* underwent adaptive evolution in strain B1 when strain A was its sole competitor. In the presence of a rich microbiota, strain B1 followed a markedly different metabolic adaptive path, with mutations in genes of D-galactonate metabolism being pervasive, an adaptive target also observed in other *E. coli* strains evolving in the gut^18,30^. Remarkably, the degree of convergent evolution between strains was substantially higher in the presence of a complex microbiota (Fig. 2b, Supplementary Figs. 7e, f). The *fim* operon and *barA* were under strong selection in both strains. The adhesin FimH, located at the tip of type I pili and involved in catch-bond formation^31^, is known to be highly polymorphic in *E. coli*^32^, and here we show that it can evolve rapidly regardless of inflammatory status, with new alleles reaching high frequencies within a strain (Supplementary Fig. 7f). The BarA–UvrY two-component signal transduction system, which plays a central role in carbon metabolism, was also under selection in the intestine. The convergent evolution observed in metabolic pathways and adhesion is consistent with adaptation to a fluctuating environment in which strains compete for overlapping niches and resources.

## Strain B1 harbours a diverse repertoire of mobilizable genomic elements

Strain B1 is known to donate DNA to strain A *via* conjugation and transduction^13,14^. To investigate the potential of its prophages for HGT, we exposed it to mitomycin C, a bacterial metabolite that induces prophage excision, and isolated the resulting phage particles^33^. Sequencing of the phage particles’ DNA revealed high coverage for nine of the ten bioinformatically predicted prophages in strain B1 (Fig. 3a, Supplementary Table 14). These include the Nef and KingRac prophages and a satellite phage, elements previously shown to transfer in the gut^14^, as well as six additional regions potentially encoding phages or phage-mobilized elements.

The data suggest a remarkable transduction potential: several prophage-adjacent genomic regions displayed moderate to high read coverage, indicating that flanking DNA may be encapsidated into viral particles (Fig. 3a). Notably, a large ∼600 kb region flanked by three prophages (Supplementary Table 14) exhibited high coverage, raising the hypothesis for lateral transduction changing the genomic structure of these *E.coli* strains^34,35^. For instance, DNA encapsidation was detected on both sides of the Nef and KingRac prophages (Fig. 3a).

Taken together, these *in vitro* findings point to a high potential for prophage transfer and prophage-driven HGT events to occur *in vivo*.

## The adaptive landscape of HGT is rich and complex under chronic inflammation

To assess the extent to which the HGT potential revealed *in vitro* is realized *in vivo*, we measured transfer frequencies throughout the colonization experiment. Using PCR-typing, we screened evolving clones of strain A for eight distinct transfer events that can occur between these strains: two conjugation events (by plasmids A and B), four intact prophages (Nef, KingRac, DMcol, JOcol), one satellite prophage (hereafter “satellite”), and one genomic region harbouring *fnr*, a gene important for gut colonization^14,36,37^.

In the absence of microbiota, transfer of plasmid A, phage Nef, and the satellite occurred at comparable frequencies in both IL10KO and WT mice (P = 0.480, P = 0.518, and P = 0.299, respectively). Other transfer events were detected only rarely, with no significant differences between host backgrounds (P = 0.138, LM) (Fig. 3b, Supplementary Table 15).

In the more ecologically relevant SPF mouse model, conjugation and transduction events were significantly elevated in the chronically inflamed gut (Fig. 3c). Plasmid B was rarely transferred in either background, whereas plasmid A transconjugants arose at significantly higher frequencies in IL10KO mice (P=0.021, day 371, LM). Prophages Nef and KingRac transferred at similar rates regardless of inflammation status (Fig. 3c, Supplementary Table 16). Importantly, the genomic region adjacent to KingRac carrying *fnr* gene, was transferred in all IL10KO mice but in only one WT mouse, strongly suggesting that chronic inflammation enhances the effective rate of transduction. Notably, strain A lysogens that acquired *fnr* reached dominant frequencies in IL10KO mice, underscoring their high selective value in the inflamed environment. Satellite transfer was likewise significantly elevated in inflamed mice (P = 0.051, LMM), with satellite-carrying strain A clones becoming dominant in all IL10KO SPF mice but in only two WT SPF mice.

Despite considerable variability across timepoints and transfer event types (Fligner-Killeen test, P=0.008), HGT frequency in chronically inflamed SPF mice was significantly higher than in their WT SPF counterparts (P=0.023, LMM) and significantly higher than in mono-colonized IL10KO mice P=0.000975, LMM). This result is consistent with the expectation that interactions between intestinal microbes trigger prophage induction^38^. Together, these results demonstrate that the presence of a complex microbiota in hosts lacking a key anti-inflammatory cytokine strongly shapes bacterial evolution through HGT.

## High rates of horizontal gene transfer promote phage diversity through prophage mosaicism

To characterize HGT events, we performed long-read sequencing on strain A and strain B1 clones isolated from faecal samples at the end of colonization (days 140 and 371) in IL10KO and WT mice (Supplementary Table 17). Long-read sequencing analysis confirmed mutations identified in short-read population sequencing (Supplementary Tables 18,19), as well as transfers identified from PCR-typing: prophages Nef, KingRac, DMcol, JOcol, and the satellite from strain B1 were detected in strain A clones (Figs. 4abc; Supplementary Table 20), alongside multiple independent transduction events containing *fnr*. A new genomic region, containing an operon known to be under selection in WT mice^39^ (*gat* operon), was found in IL10KO mouse (Figure 4bd, Supplementary Table 20).

**Figure 4.**
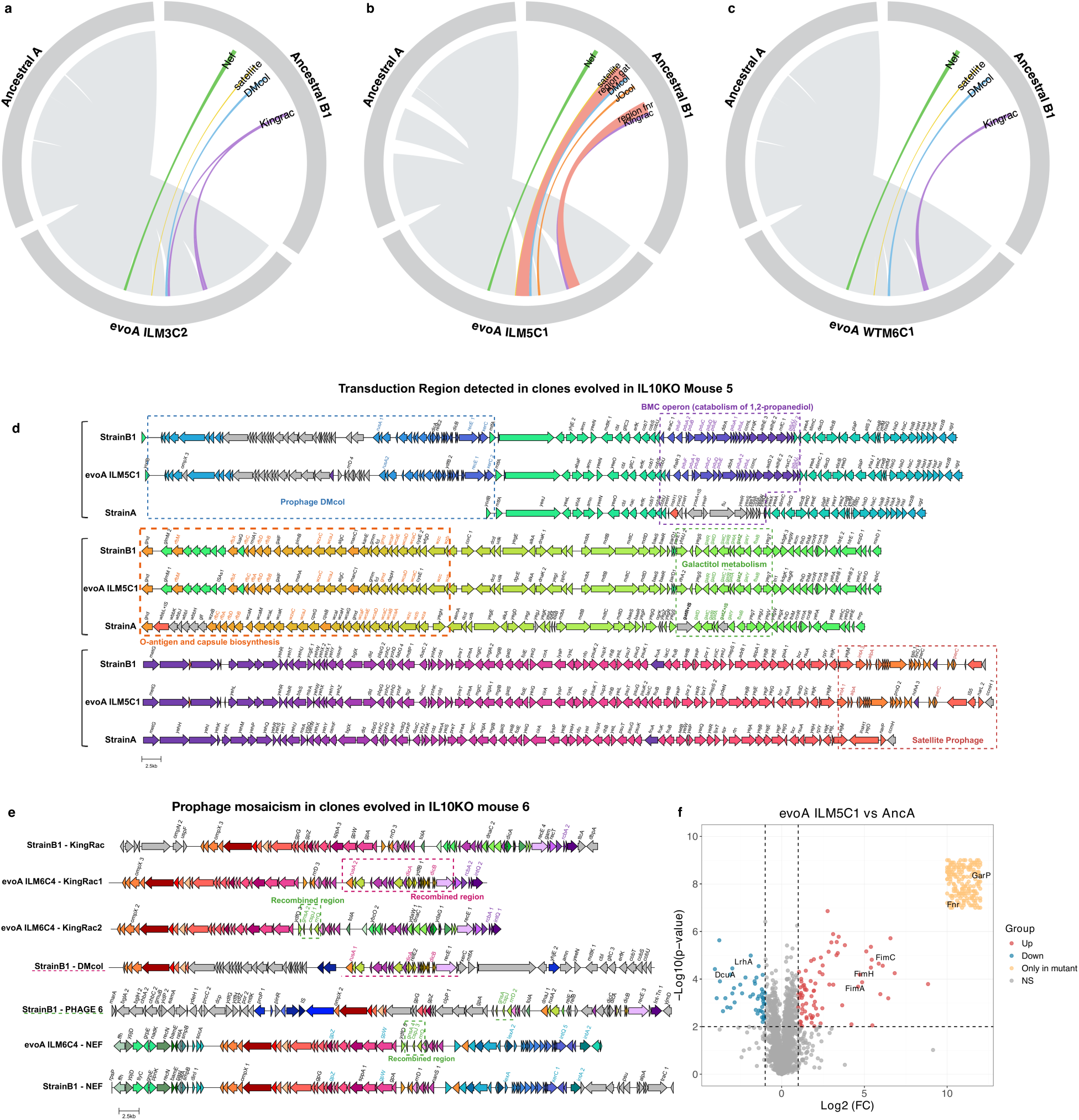
Comparison of evolved clones with their ancestors reveals hybridization. **a,** Clone evolved in IL10KO mouse 3 that acquired prophages Nef and DMcol, 3 copies of KingRac and the satellite **b,** Clone evolved in IL10KO mouse 5 that acquired two chromosomal regions of strain B1 (carrying genes (*fnr* and *gat*) under strong selection in the gut), as well as prophages Nef, DMcol, JOcol, KingRac and the satellite. **c,** Clone evolved in WT mouse 6 that acquired prophages Nef and DMcol, 2 copies of KingRac and the satellite. **d,** Hybrid clone evolved in IL10KO mouse 5 that acquired a large transduction region (248 kb) with new/restored functions: propanediol catabolism (purple), galactitol metabolism (green) and altered O-antigen (light orange). This region is flanked by prophage DMcol (blue) and satellite (orange). **e,** Hybrid clone exhibiting extensive phage mosaicism. Recombination between prophages KingRac and DMcol highlighted in pink, KingRac and prophage 6 in green and Nef and prophage 6 in green. **f,** Hybrid clone evolved in IL10KO (evoA ILM5C1) with large transductions (in panel b) shows a significantly changed proteome: proteins of type I fimbriae have higher concentration in the evolved clone compared with its ancestor, as well as proteins of colanic acid capsule, the oxygen sensor Fnr and the transporter of galactarate GarP; DcuA a C4-dicarboxylate major transporter for L-aspartate uptake under aerobic conditions and LhrA known to regulate the transcription of genes involved in the synthesis of type 1 fimbriae as well as directly controlling the expression of the master regulator FlhDC, are both at lower concentration in the evolved clone (Supplementary table 24). Highlighted in red are proteins with higher concentration in evolved IL10KO mouse 5 clone 1, in blue the proteins with lower concentration and in orange the proteins only detected in the evolved clone (including proteins encoded in the acquired plasmid).

Comparison of ancestral strain B1 prophages with those acquired by evolved strain A clones revealed extensive inter-prophage recombination. Recombination between KingRac and DMcol, JOcol, and Phage 6 (Fig.4e, Supplementary Fig.9, Supplementary Table 20), and between Nef and prophage 6 (Fig.4e, Supplementary Table 20), led to a high degree of mosaicism and genomic plasticity caused by this evolutionary process. This is consistent with the shared modular architecture between strain B1 and strain A prophages, including structural modules (tail assembly, fibronectin, and minor tail genes) and a recombination module (*recE*, *recT*, *gam*, and a viral-like recombinase). Recombination was predominant in KingRac prophage (Supplementary Fig.8, Supplementary Table 20), and surprisingly some evolved clones of strain A acquired more than one KingRac from strain B1 (Fig. 4e). Interestingly, all strain A clones that received DMcol, carried structural modules from KingRac and Nef rather than DMcol strain B1 original prophage. Recombination in A-native cryptic prophages (DLP12 and Qin) was less frequent (Supplementary Table 20). At the same time, structural variation was also detected in strain B1 evolved clones (Supplementary Table 21). Two big deletions were detected in the majority of clones, loss of prophage 6 in two clones of IL10KO mouse 5 and loss of prophage 9 in a different clone from the same mouse. Importantly, phage mosaicism was also found in strain B1 (Supplementary Table 21), showing that diversification of temperate phages is not strain dependent. Long-read sequencing confirmed the lower frequency of HGT in the absence of microbiota but also revealed a unique new event in one IL10KO mouse (Supplementary Table 22).

## *E. coli* extreme makeover: adaptation by HGT in the inflamed gut

Transduction events in strain A were both frequent and of striking functional consequence, particularly under inflammatory conditions. Beyond the acquisition of *fnr*, occurring in all SPF IL10KO mice *via* transduction spanning 22-438kb, in two IL10KO mice (M2 and M4), transduction extended to both flanks of KingRac, encompassing upstream (123-438kb) and downstream (12-43kb) regions.

A remarkable large transduction event (248 kb), flanked by DMcol and the satellite integration region, was identified in IL10KO mouse 5 at late timepoints (days 307 and 371) (Fig. 4d). This region encodes several functions, including O-antigen biosynthesis (*rfb* and *wbb* operons), causing a serotype change (Supplementary Table 23), and restoration of galactitol metabolism, as confirmed phenotypically (Supplementary Fig. 10a). Strikingly, multilocus sequence type analysis^40^ assigned the evolved lysogens, from IL10KO mice M2 and M5, to a new lineage (ST2055), distinct from that of its ancestors: strain A (ST10) and strain B1 (ST602). Massive changes caused by horizontal gene transfer, as this one, may explain the emergence of recent pandemic fluoroquinolones resistant *E. coli* strains^41^. Phylogenetic analysis confirmed that this evolutionary split was driven by HGT (Supplementary Fig. 10b). When we examined the protein composition of the hybrids, the differences were striking, with proteomics characterization revealing hundreds of proteins with altered abundance (Fig. 4f, Supplementary Fig. 10c, Supplementary Tables 24 and 25). Clone 1 from IL10KO mouse 5 (Fig. 4b, d) alone showed 334 proteins at significantly higher concentrations and 201 at lower concentrations, while clone 14 from IL10KO mouse 4 displayed an even more dramatic shift, with 1,051 proteins increased and 180 decreased.

Interestingly, both clones showed elevated levels of proteins involved in type-1 fimbriae biosynthesis, including FimH and FimC. At the same time, other proteins were reduced, LrhA, which regulates type-1 fimbriae and flagellar synthesis, was diminished in the clone from mouse 5, and OmpC, a key porin linked to azithromycin and colistin susceptibility³⁸, was reduced in the clone from mouse 2. Together, these results show that recombination, amplified in the conditions of a chronic inflamed gut, can drive sufficiently large genomic change to alter the evolutionary identity of a bacterial lineage within a single host.

## Discussion

Millions of patients suffer from IBD^42^, and the ecology of their gut microbiomes is changed under such disease conditions^43^. A better understanding of the mechanisms underlying IBD is urgently needed in face of the rise of this disease across the world^42^. Here, using experimental evolution in mouse models of IBD with different microbiome compositions, we show that the *tempo* and *mode* of *E. coli* evolution is significantly altered under chronic inflammation. As observed in humans^44^, the abundance of commensal *E. coli* is higher in the inflamed gut of IL10KO mice with a complex microbiome. However, when *E. coli* is the sole colonizer of IL10KO mice it reaches the same abundance as in WT mice, suggesting that the boosting of *E. coli* in an inflammatory environment depends on the presence of other species. Importantly, the rate of mutation accumulation in strains of *E. coli* is boosted when they colonize IL10KO mice, and several genes of *E. coli* adaptation are diagnostic of this disease condition. Notably, *E. coli* adapts to the specific metabolic conditions of the inflamed gut by mutations in genes encoding glycerol (*glpR*) and hexuronates (*exuT*) metabolism as well as glycerate, glycolate, glucarate and galactarate metabolism (*garK*, *garL*, *gudP*, *cdaR*). Remarkably, the *gar/gud* pathway that converts oxidized sugars, underwent convergent evolution across different species of commensals enriched in the inflamed gut^45^. Consistent with the increased oxidative stress under inflammation^46^, *E. coli* repeatedly evolves a non-synonymous substitution (N22S) in the stringent starvation response gene *sspA* in all IL10KO but in none of the WT mice studied. Furthermore, evolution in genes with cell-wall functions, including the outer-membrane (*waa* operon) and capsule formation (*yjjQ*, *gfcA*, *etk*), were specific to IL10KO mouse adaptation, suggesting that the interface between bacteria and host is under selection in this environment.

Beyond mutation we find phage-driven HGT between *E. coli* strains to have a remarkable power at accelerating their evolution in the inflamed gut. Our data shows extensive bacterial DNA transferring between strains, together with high levels of recombination between their integrated phages (phage mosaicism^47^), suggesting that high rates of prophage excision are occurring in the inflamed gut. Recently increased abundance of temperate phages in patients with Crohn’s disease, a form of IBD^48^, in comparison to healthy controls has been reported. Our data in IL10KO vs WT mice, strongly suggests that the incorporation of both *E. coli* genomics and transductomics^49^ of its prophages may constitute a useful predictor of these disease conditions. Considering that prophages can transfer virulence genes, the high rates of prophage movement and mosaicism observed, suggest that the inflamed gut constitutes a unique environment for fostering the spread and evolution of pathogenic traits. At the same time, the increased plasmid transfer observed under chronic inflammation also suggests increased rates of antimicrobial resistance in IBD patients. Overall, the results of evolution in mice presented here make two testable predictions with important consequences for human health and microbial evolution: *i*) genomic changes in *E. coli*, a human commensal, may constitute a new biomarker of IBD; *ii*) given that the number of individuals afflicted with inflammatory diseases is in the millions and has been increasing, it is expected that an acceleration of *E. coli* molecular evolution is occurring in these hosts.

## Material and Methods

### Animals and Ethics

C57Bl/6J mice (six to eight-week-old) were bred by the Rodent facility at Gulbenkian Institute for Molecular Medicine (GIMM) (Oeiras, Portugal). A total of 24 mice were used: 6 wild-type Germ-free (WT-GF) mice and 6 IL10KO (model for inflammatory bowel diseases - IL10-/-, or IL10KO-GF) mice (males and females) and 6 wild-type specific-pathogen free (WT-SPF) mice and 6 IL10KO (model for inflammatory bowel diseases - IL10-/-, or IL10KO-SPF) mice (females). Food and drinking water were provided *ad libitum*. All mice were housed in ventilated specific-pathogen free conditions and in an animal facility in accordance with GIMM guidelines and European recommendations. All animal experiments were approved by Ethics Committee of GIMM (license reference: A006.2023), and by the Portuguese National Entity that regulates the use of laboratory animals (DGAV - Direcção Geral de Alimentação e Veterinária (license reference: 74842/25-S). All experiments conducted on animals followed the Portuguese (Decreto-Lei n° 113/2013) and European (Directive 2010/63/EU) legislations, concerning housing, husbandry and animal welfare.

### Bacterial Strains

*E. coli* strains used in this study express fluorescent proteins and carry antibiotic resistance markers. *E. coli* strain A expresses either a Yellow or a Cyan Fluorescent Protein, namely (YFP) or (CFP), respectively, together with an ampicillin (YFP) or chloramphenicol (CFP) resistance markers, as well as a streptomycin resistance mutation. The strains are MG1655 DM08-YFP gatZ::IS1 [galK::amp (pZ12)::PLlacO-1-YFP, strR (rpsL150), ΔlacIZYA::scar] and MG1655 DM08-CFP gatZ::IS1 [galK::cm (pZ12)::PLlacO-1-CFP, strR (rpsLl50), ΔlacIZYA::scar]. *E. coli* strain B1, a mouse commensal, expresses a Red (mCherry) or a Green Fluorescent Protein (sfGFP) and a chloramphenicol resistance marker. *E. coli* clones from both strains were grown at 37°C under aeration in liquid media Lysogeny Broth (LB Medium AppliChem A0954) and LB agar plates (VWR 84684.0500). Media were supplemented with antibiotics streptomycin (100 µg/mL), ampicillin (100 µg/mL) or chloramphenicol (30 µg/mL) when specified. Serial plating of 1X PBS dilutions of faeces in LB agar plates supplemented with the appropriate antibiotics were incubated overnight and YFP, CFP, mCherry and sfGFP-labeled bacterial numbers were assessed by counting the fluorescent colonies using a fluorescent stereoscope (Zeiss Stereo Lumar V12). The detection limit for bacterial plating was ∼300 CFU/g of feces^18^.

### *In vivo* Coevolution Experiment

All specific-pathogen free (SPF) C57BL/6J mice WT and IL10KO were given a streptomycin (5 g/L) treatment in the drinking water for 7 days, to break colonization resistance, followed by 2 days in which streptomycin treatment was absent to clean the gut from antibiotic traces. During this time, mice were kept co-housed to homogenize the mouse microbiota.

The colonization inoculum was prepared by inoculating the four bacterial clones (two colour versions of the two strains) individually from frozen stocks in 5 ml of Brain Heart Infusion broth supplemented with the appropriate antibiotics (streptomycin (100 µg/mL) for the A strain or chloramphenicol (30 µg/mL) for the B1 strain), and let them grow overnight with aeration at 37°C. The next day, we inoculated 20 mL of the first growth solution into 2 mL of BHI without antibiotics and adjusted bacterial density (OD600=2). We then prepared a bacterial suspension in 1X phosphate buffered saline (PBS) solution approximately 1.6x10^8^ CFUs containing the four clones at equal quantities (1:1:1:1 mixture). The animals, including the SPF and GF, both IL10KO and WT, were inoculated by intragastric gavage with 100 µL suspension of a mixture of both *E. coli* strains. All mice were individually housed, SPF in ventilated cages in SPF barrier conditions and GF mice were kept isocages (Tecniplast) at the GIMM rodent facility. Faecal pellets were collected during approximately 5 months from GF mice and for approximately 1 year (371 days) in SPF mice and stored in 15% glycerol at -80°C for later analysis.

### Faecal Lipocalin-2 quantification

To measure Lipocalin in the faeces, we used the faecal samples obtained from the mice that were frozen at -80 °C. The faecal pellets were homogenized, vortex for 5 minutes, and centrifuged for 15 minutes at 18000 g, 4°C. Lipocalin levels were estimated in the supernatants using murine Lipocalin ELISA kit (Mouse Lipocalin-2/NGAL ELISA, R&D Systems) according to manufacturer’s instructions.

### IgA quantification assay

Total faecal IgA was measured by sandwich ELISA. For detection of total IgA, ELISA plates (Nunc) were coated with 5 µg/ml of goat anti-mouse IgA (Southern Biotech) capture antibody and incubated overnight at 4°C. Plates were washed and blocked with 1% BSA in PBS for 1h at room temperature. Diluted samples and standard were added, and plates incubated for 2h at room temperature. Captured IgA was detected by horseradish peroxidase (HRP)-conjugated goat anti-mouse IgA antibody (Southern Biotech). ELISA plates were developed by TMB microwell peroxidase substrate (R&D Systems) and quenched by 1M H2SO4. Colorimetric reaction was measure at OD = 450 nm by a Microplate Reader.

### Populations sequencing and analysis

Evolved clones from GF WT mice were sequenced in Ameline et al 2025^24^. Evolved clones from WT SPF mice at days 43 and 88 were sequenced in Frazão et al 2025^14^. For the other mice and timepoints, DNA was extracted with Phenol-Chloroform from a pool of *E. coli* clones (∼1000) after plating faecal samples taken from each mouse on LB plates with antibiotic specific for each strain. DNA concentration and purity were quantified using Qubit and NanoDrop, respectively.

For GF IL10KO evolved populations the DNA library construction and sequencing were carried out by the GIMM genomics platform using the Illumina Nextseq2000 platform for samples of day 28 and day 140. For days 84 and 111 DNA library construction and sequencing were carried out by Novogene using the Illumina NovaSeq X Plus Series (PE150) platform.

For SPF evolved populations the DNA library construction and sequencing were carried out by the GIMM genomics platform using the Illumina Nextseq2000 platform for days 151, 371 from WT mice and days 43, 88, 244 from IL10KO mice. For day 244 from WT mice and days 151 and 371 from IL10KO mice DNA library construction and sequencing were carried out by Novogene services using the Illumina NovaSeq X Plus Series (PE150) platform. Raw sequencing reads were processed using fastp^50^ (version 0.23.2). Sequencing adapters were removed, and raw reads were trimmed bidirectionally by 4bp window sizes, retaining an average base quality of 20. Reads shorter than 100 bp and those with more than 50% of bases having a Phred score below 20 were removed, followed by base correction of overlapping reads. Moreover, duplicated reads were also removed, using dedup function of fastp. References genomes for alignment of sequenced reads were K-12 substrain MG1655; Accession Number: NC_000913.2, for the alignment of reads from strain A; Accession Number: SAMN15163749 for the alignment of reads from strain B1; Accessions CP054663 and CP054664 for the two plasmids of strain B1 (with length 108557 bp and 68935 bp). BBsplit (version 38.90) (https://www.osti.gov/servlets/purl/1241166) was used to remove reads with sequences highly divergent from the reference genome. Variant calling in the evolved populations was performed using the 0.37.1 version of the BRESEQ pipeline^51^ with the polymorphism option on and default settings, except for: a) polymorphism minimum variant coverage of 5 reads; b) base quality cutoff of 30; c) minimum mapping quality of 20. To decrease the probability of detecting false positives in the evolved populations, sequencing reads from the ancestral clones were used as controls during variant calling. Variants predicted in the ancestral clones using the BRESEQ pipeline in polymorphism mode (using the parameters described above) were considered as false positives and excluded from the evolved population analysis. Only mutations >= 10% frequency in the population were considered for the analysis. A previously designed custom R script was used for identification of two large deletions detected in strain B1^14^(available at https://github.com/hugocbarreto/Lateral-gene-transfer-causes-genomic-repair-when-strains-coexist-in-the-gut).

Variant calling on some samples from SPF mice for strain A and strain B1 revealed thousands of variants across the reference genome. Using BLASTN^52^, we identified reads with 100% identity with genomes from *Enterobacter hormachei* and *Leclercia adecarboxylata*. For these samples, BBSplit was run including genomes from *E. hormachei* (Accession Numbers: CP010376.2, CP019889.1, CP021137.1, CP031565.1, CP099314.1, CP099316.1, CP132232.1, CP141537.1, LR607340.1, LT840187.1) and *L. adecarboxylata* (CP106959.1, LR590464.1). These genomes were selected based on a 100% match of the contaminant reads with NCBI database. BBSplit was run including the reference genome of strain A (K-12 substrain MG1655; Accession Number: NC_000913.2) or strain B1 (from Accession Number: SAMN15163749 for the alignment of reads from strain B1). Then, all samples were re-analysed using the BRESEQ pipeline as described above.

We have previously shown that certain horizontal gene transfer (HGT) events can occur between strain B1 and strain A^14^. Specifically, the transfer of different prophages and transductions from strain B1 to strain A can be detected using the BRESEQ pipeline, as hundreds of variants confined to certain genomic regions in strain A. Importantly, the reads of these variants match 100% with the genome of strain B1, supporting HGT from strain B1 to strain A. The presence and frequencies of these events were verified by PCR typing (see methods below).

### PCR-typing for plasmids, prophages and transductions events

PCR-typing for the two plasmids and phage Nef and KingRac, as well as the *fnr* region was performed as previously^14^ to determine the frequency of lysogens in the strain A *E. coli* populations.

For the remaining phages specific primers were design for each region:

satellite_Forward – GCACACAATGACGGGCTTTT

satellite_Reverse - TGTCCTGATTTGCCATCCCC

DMcol_Forward – AGGGGAGCAGAGTAAAACGC

DMcol_Reverse - TAGCACTTGCCCTTCGGTTT

JOcol_Forward – TCCAGTCACGCCAACAAGAG

JOcol_Reverse - GTAGACCAGCGACAGGAAGG

PCR amplification was performed in randomly isolated 20-24 clones from strain A evolved *E. coli* populations. PCR reactions were performed in a total volume of 25 μL, containing 1 μL of each clone growth in liquid LB media, 1X Taq polymerase buffer, 200 μM dNTPs (GRISP), 0.2 μM of each primer and 1.25 U Taq polymerase (Thermo Scientific). PCR reaction conditions: 95°C for 3 min, followed by 35 cycles of 95°C for 30 s, (depending region)°C for 30 s and 72°C for 30 s, finalizing with 5 min at 72°C. DNA was visualized on a 2% agarose gel stained with Xpert Green (GRISP).

### Metabolite Identification Using ¹H-NMR

Faecal samples of SPF mice from the day 57 were prepared for proton Nuclear Magnetic Resonance (¹H-NMR) analysis by adding 1 mL of deuterated water (D₂O, 99.9%, Sigma-Aldrich) and 0.3 mg of 1 mm glass beads (Scientific Industries, SI-BG01). Samples were homogenized using a QIAGEN Retsch Tissuelyser II with two 1-minute bead-beating cycles at 30 Hz/s with plate inversion between cycles. The homogenates were then centrifuged at 14,000 RPM for 30 minutes at 4°C. The supernatant was filtered through a 0.22 μm syringe filter (Avantor) and then through a 3 kDa filters (Vivaspin® 500) at 15,000 RCF for 6 hours at 4°C.

For NMR analysis, a mixture of 150 μL of the filtered was mixed with 380 μL of D₂O, 60 μL of phosphate buffer (pH 7.10 and with 2% sodium azide), and 10 μL of 0.05% (w/v) TSP-d₄ (3-(trimethylsilyl)propionic acid-d₄ sodium salt, Sigma-Aldrich, 0.04949 mM) as a chemical shift reference. This was prepared into a 5 mm NMR tube, homogenized by inversion and the pH was measured.

Spectra were acquired using a Bruker AVANCE II+ 500 MHz NMR spectrometer equipped with a 5 mm TCI(F)-z H-C/N Prodigy cryoprobe. A 1D NOESY pulse sequence with pre-saturation (noesypr1d) was used, with a calibrated 90° pulse and 100 ms of mixing time. The acquired spectrum was 200 scans, with an acquisition time of 4 s, and 1 s relaxation delay, using 48k data points and a spectral width of 6002 Hz (12ppm), at 25°C.

Phase correction and spectral processing were performed, and metabolites were identified manually using the Chenomx NMR Suite (v8.11). Final metabolite concentrations were normalized based on dilution, filtration, and sample weight.

For statistical analysis, the processed data were imported into R (v4.4.2) via RStudio (v2024.12.1.563). A Wilcoxon test was applied to each metabolite, followed by false discovery rate (FDR) correction. A volcano plot was then constructed using the FDR-adjusted p-values, applying a significance threshold of FDR < 0.05 and a fold change (FC) cutoff of 2, with FC based on median concentration.

### Extraction, sequencing, and analysis of DNA encapsidated in the phage head (capsid)

A total of 70 mL of the phage lysate from *E. coli* strain B1 was produced by mitomycin C induction. After centrifugation (4,000 rpm, 15 min, 4°C) and filtration (500-mL filtropur BT50 top filter, 0.2 μm), the phage lysate was treated with RNase (1 μg/mL) and DNase (10 μg/mL) for 30 min at room temperature, before addition of 1 M NaCl for 1 h on ice. The lysate solution containing the induced phages was loaded onto a sucrose solution (30%) and ultracentrifuged at 80,000 g for 2 h at 4°C. After ultracentrifugation, phage lysates were treated for 30 min with DNase (10 μg/mL) at room temperature, with DNase being inactivated with 5 mM EDTA for 10 min at 70°C. Subsequently, an equal volume of a lysis mix (2% SDS and 90 μg/mL proteinase K) was combined with the phage lysate and incubated at 55°C for 1 h. Encapsidated phage DNA extraction was performed with an equal volume of phenol:chloroform:isoamyl alcohol solution (25:24:1). Samples were then centrifuged at 12,000 g for 5 min and the DNA obtained from the aqueous phase. Phage DNA was precipitated with ethanol (100%) and NaAc (3 M, pH 5), with the resulting DNA being resuspended in 100 μL of Tris-HCl solution (10 mM, pH 8.5)^13,53^. Encapsidated DNA was quantified using Qubit™ dsDNA High Sensitivity (Thermo Fisher Scientific). Library preparation, including DNA tagmentation, PCR-mediated adaptor addition and amplification of the adapted libraries, was performed following the Nextera XT library preparation protocol (Illumina), as previously described^54^. Libraries were confirmed using Fragment Analyzer (Agilent Technologies) and sequencing performed using the Illumina NextSeq 2000, obtaining 2 × 65 bp pair-end reads. Library preparation and sequencing were performed by the GIMM Genomics Platform. Approximately 400 M reads were generated. The reads where then mapped to the reference genomes of strain B1 (Accession Number: SAMN15163749) using BBmap, version 39.01(https://packages.guix.gnu.org/packages/bbmap/39.01/) with default parameters with the following alterations: ambiguous=random, qtrim=lr, minid=0.97^55^. The resulting read mapping files were sorted and indexed using SAMtools (version 1.18), followed by conversion to bigwig using deepTools bamCoverage with the following parameters: --binSize = 100^56,57^. The R packages ggplot2 3.4.452 and ggcoverage (1.2.0) were used for creating Fig.3a^58,59^.

Prophage regions were predicted using Phastest^60^ (https://phastest.ca/).

### Nanopore sequencing and analysis

DNA extraction was performed using the Monarch HMW DNA Extraction Kit for Tissue (New England Biolabs -T3060L) from several clones from both strain A and strain B1 isolated at the last timepoint of colonization in both IL10KO and WT mice (d371, with exception for strain A in WTM1 and M2 which clones were isolated at d151 and d244 respectively). DNA concentration and purity were quantified using Qubit and NanoDrop, respectively. A clean-up pre-library was performed using 0.7x AMPure beads (Beckman Coulter A63880). Libraries were prepared using Rapid Barcoding and Native Barcoding Kits from Oxford Nanopore (details in Supplementary Table S19).

Raw nanopore reads were pre-processed prior to assembly. Untrimmed external adapters and reads containing potential internal adapters and chimeras were identified and removed using Porechop (https://github.com/rrwick/porechop). Quality filtering was subsequently performed with Filtlong (https://github.com/rrwick/filtlong), retaining reads with a minimum length of 1,000 bp and a minimum mean quality score of Q10. For more complex genomes, a subset of ultra-long reads (over 20000bp) was done to resolve genome continuity. Per-barcode read quality metrics were assessed using NanoPlot (https://github.com/wdecoster/nanoplot). De novo genome assembly was carried out using Flye^61^ (v.2.9.6), using a subsampled coverage of 20× to reduce computational load and ensure robust base frame for genome assembly. Draft assemblies were polished using Medaka^©^ (v2.2.1): reads were first aligned to the draft assembly using mini_align, and consensus inference was then performed with Medaka’s inference module across all contigs simultaneously, followed by sequence stitching to produce the final polished assembly. Structural annotation of the polished assemblies was performed with Prokka^62^ (v1.14.6), specifying *Escherichia coli* as the target organism and using the --compliant and --metagenome flags.

Following annotation, each clone’s genomic contig was aligned to the ancestral reference genome in Mauve to assess orientation. Contigs assembled in the reverse complement orientation relative to the ancestral genome were reverse-complemented prior to downstream analysis. To ensure consistent coordinate systems across clones, all assemblies were rotated to begin at the equivalent start position of the ancestral genome, using a custom script that aligns the clone sequence to a 150-200 bp window extracted from the ancestral genome origin. Rotated assemblies were re-annotated with Prokka^62^ using identical parameters. Comparative genomics was performed by aligning the rotated annotated assemblies to the ancestral genome in Mauve^63^, and genomic alterations were manually identified and extracted at nucleotide resolution with their corresponding coordinates recorded in Geneious. In parallel, short variants and structural variants were called from the polished reads using Clair3 and Sniffles2, respectively. Final outputs, including rotated assemblies, annotations, variant calls, and manually extracted altered sequences, were consolidated for downstream analysis. For prophage recombination verification and visualization we used clinker (https://github.com/gamcil/clinker)^64^.

### Galactitol metabolism recovering phenotype

In certain *E. coli* evolved clones from strain A isolated from IL10KO mouse 5, the transduction of a region including the *gat* operon, responsible for the metabolism of galactitol, was detected. Ancestral strain A lacks the ability to metabolize this sugar-alcohol. Thus, to confirm the evolved clones gain of function we performed a phenotype assay, as previously described^22^. Briefly, we grew each clone in MacConkey plates supplemented with 1% galactitol^22^. Plates were grown at 30°C for 24 to 48 hours. Phenotypes were compared with strain A ancestral, not able to metabolize galactitol, and strain B1, which is able to metabolize it. Phenotypes are red or white colonies (consuming or not the galactitol, respectively).

### Serotype prediction

Some strain A evolved clones, isolated specifically from IL10KO mouse 5, show the acquisition of a genomic region that include several operons involved in O-antigen biosynthesis, *rfb*, *wca*, *wzc*, *wza*. These changes suggest an alteration in serotype. Using ECTyper^65^ (https://github.com/phac-nml/ecoli_serotyping) we performed an *in silico Escherichia coli* serotyping and species prediction for the clones acquiring the region and compared it with both ancestors (strain A and B1) and with clones isolated from other IL10KO and WT clones.

### Phylogenetic Analysis

Whole-genome sequences of evolved *E. coli* clones obtained from long-read sequencing, isolated throughout the co-colonization experiments, were aligned against the reference genome NC_000913.2 (strain A) using Snippy, with variants called independently for each sample using 8 CPUs. A core SNP alignment was generated across all samples using snippy-core, producing a full alignment of 4,639,675 nucleotide sites spanning 32 sequences, of which 73,662 sites were parsimony-informative (1.95% of all sites) and 98.05% were invariant. As all sequenced isolates are experimentally evolved clones derived from defined ancestral strains within a controlled *in vivo* setting, recombination filtering was not applied prior to phylogenetic inference. Maximum likelihood phylogenetic reconstruction was performed using IQ-TREE^66^ (v3.0.1). The best-fit substitution model was selected using ModelFinder^67^ under the Bayesian Information Criterion (BIC), which identified TVM+F+I as the optimal model. This model incorporates transversion-specific rate parameters with empirical base frequencies and a proportion of invariable sites (estimated at 96.6%). Branch support was assessed using both the ultrafast bootstrap approximation (UFBoot^68^, 1000 replicates) and the SH-aLRT test (1000 replicates). The tree is presented rooted at mid-point. Branches with ultrafast bootstrap support ≥ 95% and SH-aLRT support ≥ 80% were considered well-supported. Visualization was performed using iTOL (https://itol.embl.de/).

### Proteomics

Sample Preparation: For bacterial pellets preparation, 100 ml from an overnight culture was inoculated in 10ml LB and grown at 37°C, shaking, until it reached an optical density (OD) between 0.5-0.6. Afterwards cells were centrifuged in 50ml tubes at 4000rpm for 10 min at 4°C, washed in PBS and centrifuged again. For the second wash the samples were passed to 2ml Eppendorf and washed in PBS again, centrifuged for 7min at 8000rpm, the supernatant was removed and cells frozen at -80°C. Cell pellets were then resuspended in lysis buffer containing 5% sodium dodecyl sulphate (SDS), 10 mM tris(2-carboxyethyl)phosphine (TCEP) and 100 mM triethylammonium bicarbonate (TEAB), followed by incubation at 95°C for 10 min. Cells were disrupted by ultrasonication using the PIXUL system (Active Motif) for 30 min with default settings (pulse 50 cycles, PRF 1 kHz, burst rate 20 Hz). Protein content was determined by a BCA assay. Sample alkylation was performed by addition of 20 mM iodoacetamide (IAA) and incubation at 25°C for 30 min with gentle shaking. 20 μg of protein was digested using the S-Trap protocol according to the manufacturer’s instructions (ProtiFi). Peptide concentration was determined using a UV-based assay (Infinite M Nano, Tecan).

LC-MS/MS Data Acquisition: Peptides were separated by reversed-phase liquid chromatography on a Vanquish Neo system (Thermo Fisher Scientific) using a 150 mm × 75 μm C18 column at a flow rate of 350 nl/min. Mobile phase A consisted of water with 0.1% formic acid and mobile phase B of 80% acetonitrile with 0.1% formic acid. Peptides were eluted over an 80 min gradient from 1% to 95% mobile phase B. Mass spectrometric analysis was performed in data-independent acquisition (DIA) mode on an Orbitrap Exploris 480 instrument (Thermo Fisher Scientific). Full MS scans were acquired at a resolution of 120,000 over a scan range of 390–910 m/z, with an AGC target of 3.0 × 10⁶ and automatic maximum injection time. DIA MS/MS scans were acquired at a resolution of 15,000 over a precursor mass range of 400–900 m/z, using 50 variable isolation windows of 10 m/z with 1 m/z overlap, HCD collision energy of 28%, and a maximum injection time of 22 ms.

Data Processing and Statistical Analysis: Raw data were imported into Spectronaut (v20.4.260109.92449, Biognosys) and analysed using the directDIA workflow. Database searches were performed against the *E. coli* K12 proteome (UniProt UP000000625), supplemented with the proteomes of the sequenced strains when relevant. Cysteine carbamidomethylation was set as a fixed modification. Methionine oxidation and N-terminal methionine excision were set as variable modifications. Results were filtered at 1% false discovery rate (FDR) at the spectrum, peptide and protein levels. Protein-level quantification was based on the log₂-transformed protein group quantity (PG.Quantity) as reported by Spectronaut, retaining only entries with a protein group q-value < 0.01. Proteins were required to be detected in at least 2 out of 3 biological replicates in at least one experimental condition. Intensities were median-normalized across samples using observed values only. Statistical testing for differential protein abundance was performed in Perseus (v2.0) using a two-sample t-test with permutation-based FDR correction (FDR = 0.05, s0 = 0.1). Proteins with |log₂FC| > log₂(2) and p < 0.05 were additionally classified as classically significant. Data analysis and visualization were performed in R (v 4.4.0) using the tidyverse, ggplot2 and ggrepel packages.

### Statistical Analysis

All statistical analyses were performed in R (v4.4.0). Linear mixed models (LMMs) were fitted using the lme4^69^ (v2.0.1) and lmerTest^70^ (v3.2-1) packages. Faecal lipocalin-2 (ng/g) and faecal IgA (µg/g) were analysed using LMMs with outcomes log-transformed prior to modelling, genetic background (WT or IL10KO) and day as fixed effects, and a per-mouse random intercept to account for the correlation structure of repeated measurements. Only post-colonisation timepoints were included in both SPF and GF experiments for both outcomes. The effect of genetic background was evaluated by likelihood ratio test (LRT), and 95% Wald confidence intervals are reported for fixed-effect estimates.

To characterise the temporal dynamics of faecal *E. coli* colonisation, LMMs were constructed with Log₁₀-transformed bacterial loads as the response variable, genetic background and day as fixed effects, and a per-mouse random intercept. The contribution of genetic background was evaluated by comparing nested models, with and without background as a fixed effect, using F-tests with Satterthwaite-approximated degrees of freedom. Models were fitted separately for SPF and GF co-colonisation experiments.

The number of mutational targets was compared between WT and IL10KO mice using the Wilcoxon rank-sum test, applied to each strain–microbiota combination separately. To test the power of mutational targets for distinguishing host cohorts, principal component analysis (PCA) was performed on the binary mutational target matrix using prcomp with unit-variance scaling, after removing columns with no variation across samples. Analyses were conducted separately for strain A and strain B1 in both SPF and GF conditions. Pairwise Jaccard dissimilarities were computed using the vegan package and used to assess differences in mutational target profiles between genetic backgrounds by PERMANOVA (adonis2, 999 permutations). Homogeneity of multivariate dispersion between groups was verified using betadisper followed by a permutation test, as an assumption check for PERMANOVA. ANOSIM (999 permutations) was additionally performed as a complementary measure of group separation. Confidence ellipses on PCA plots assume multivariate normality.

Mutation accumulation over time was described by fitting the sum of mutation frequencies M(t) as a function of day using nonlinear least squares. For strain A in both SPF and GF conditions, and for strain B1 in SPF conditions, a linear model M(Day) = *a* × Day was used, where the rate parameter a quantifies the per-day mutation accumulation rate. For strain B1 in GF conditions, a hyperbolic model M(Day) = *a* × Day / (*b* + Day) provided superior fit, as determined by R² and AIC comparison. Models were fitted independently for each genetic background, strain, and microbiota condition, and rate parameters are reported with standard errors and t-statistics.

Horizontal gene transfer (HGT) frequency was modelled using LMMs with genetic background and day as fixed effects and mouse as random effect, fitted separately for SPF and GF datasets and for each HGT region type. Background differences in HGT frequency were additionally examined at each timepoint independently using simple linear models. Global HGT occurrence, defined as a binary outcome (frequency > 0) aggregated per mouse across all timepoints, was compared between backgrounds by linear regression, with a separate model additionally adjusting for day. The intensity of HGT events, quantified as continuous frequency values across all observations, was modelled using an LMM with background and day as fixed effects and mouse as random effect. Homogeneity of variance between backgrounds was assessed using the Fligner-Killeen test. To determine whether microbiota complexity influenced HGT occurrence, SPF and GF data from IL10KO mice were combined and analysed using a generalised linear mixed model with binomial response, environment as a fixed effect, and random intercepts for both mouse and day. A P-value threshold of 0.05 was applied throughout. For statistical analysis of metabolomics and proteomics data see the specific respective methods sections.

## Data Availability

Sequencing data is available from the NCBI SRA database under the Bioproject accession number PRJNA1272826. All other relevant data are within the paper and its supplementary data.

## Acknowledgments

We thank Camille Ameline and Daniela Guleresi for help with the experiments of germ-free mice, Beatriz Mindrico for help in R scripts for the PCA analysis, Rui Meneses and João Costa (GIMM Genomics facility) for help in setting up the nanopore technology and its analysis pipeline in the lab. We thank all the lab members of Evolutionary Biology lab for valuable discussions.

## Author contributions

E.S., N.F., M.L. and I.G. conceived and designed the study. E.S. and N.F. performed in vivo colonisation experiments and followed *E. coli* abundances and lipocalin-2 and IgA measurements. M.L., E.S., P.M, N.F. performed *in vitro* experiments, N.F. performed phage induction, purification, sequencing and analysis of DNA inside phage capsids, P.M., M.L. performed DNA extraction, sequencing and analysis of nanopore sequenced clones, M.L., I.G. performed genomic comparative analysis of evolved populations, P.Mo., K.X. performed and analysed metabolomics experiments, M.A., M.L., I.G. performed proteomics experiments and analysis. M.L. and I.G. led final data analysis, data organization and repository preparation. Funding was acquired by I.G. and K.X. The manuscript was drafted and finalised and all figures were prepared by M.L. and I.G. All authors approved the final version of the manuscript.

