## Supplementary figures for "Pervasive bacterial and prophage hybridization during chronic gut inflammation"

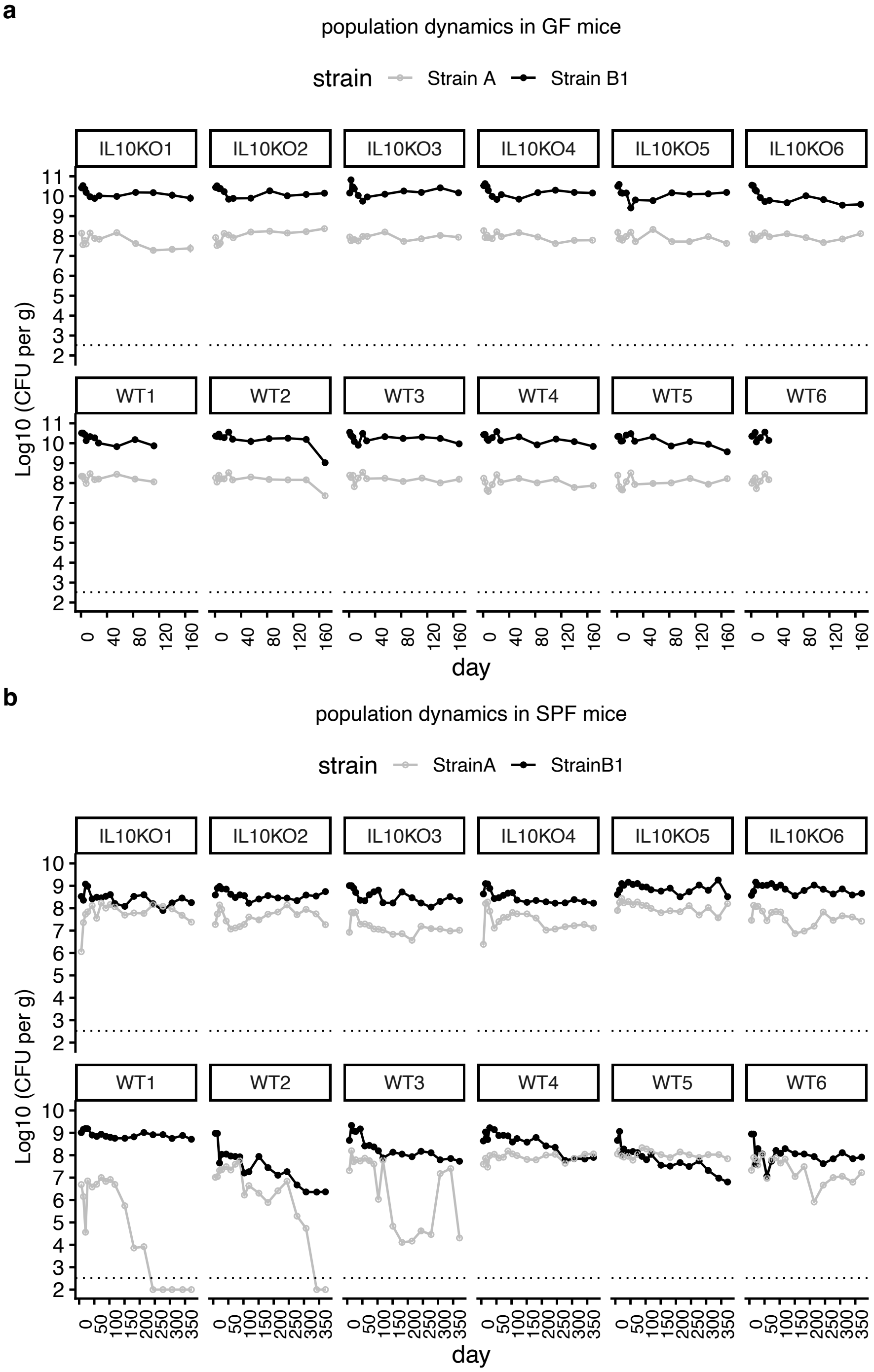


**Supplementary Figure 1. Time series of relative abundances of the two *E. coli* strains**. Strain A colonizes with lower abundance than strain B1 both mouse genotypes in (**a**) the absence of microbiota (germ-free) with IL10KO mean $Log_{10}$CFUs of strain A: 7.89 ($\pm$0.08, sem.) *vs*. mean $Log_{10}$CFUs of strain B1: 10.09 ($\pm$0.11, sem) $g^{-1}$ of feces (T-test, Pvalue= $1.98x10^{-11}$) and WT mean $Log_{10}$CFUs of strain A: 8.10 ($\pm$0.10, sem) vs. mean $Log_{10}$CFUs of strain B1: 10.14 (+0.03, sem) $g^{-1}$ of feces (T-test,Pvalue= $2.6x10^{-12}$). **b**, similar pattern seen in the presence of microbiota, with IL10KO mean $Log_{10}$CFUs of strain A: 7.6 ($\pm$0.24, sem.) *vs*. mean $Log_{10}$CFUs of strain B1: 8.64 ($\pm$0.16, sem) $g^{-1}$ of feces (T-test, P= $3.1x10^{-5}$). WT mean $Log_{10}$CFUs of strain A: 7.08 ($\pm$0.65, sem) vs. mean $Log_{10}$CFUs of strain B1: 8.21 ($\pm$0.41, sem) $g^{-1}$ of feces (T-test, P= 0.016). (Supplementary Tables 5,6 respectively).

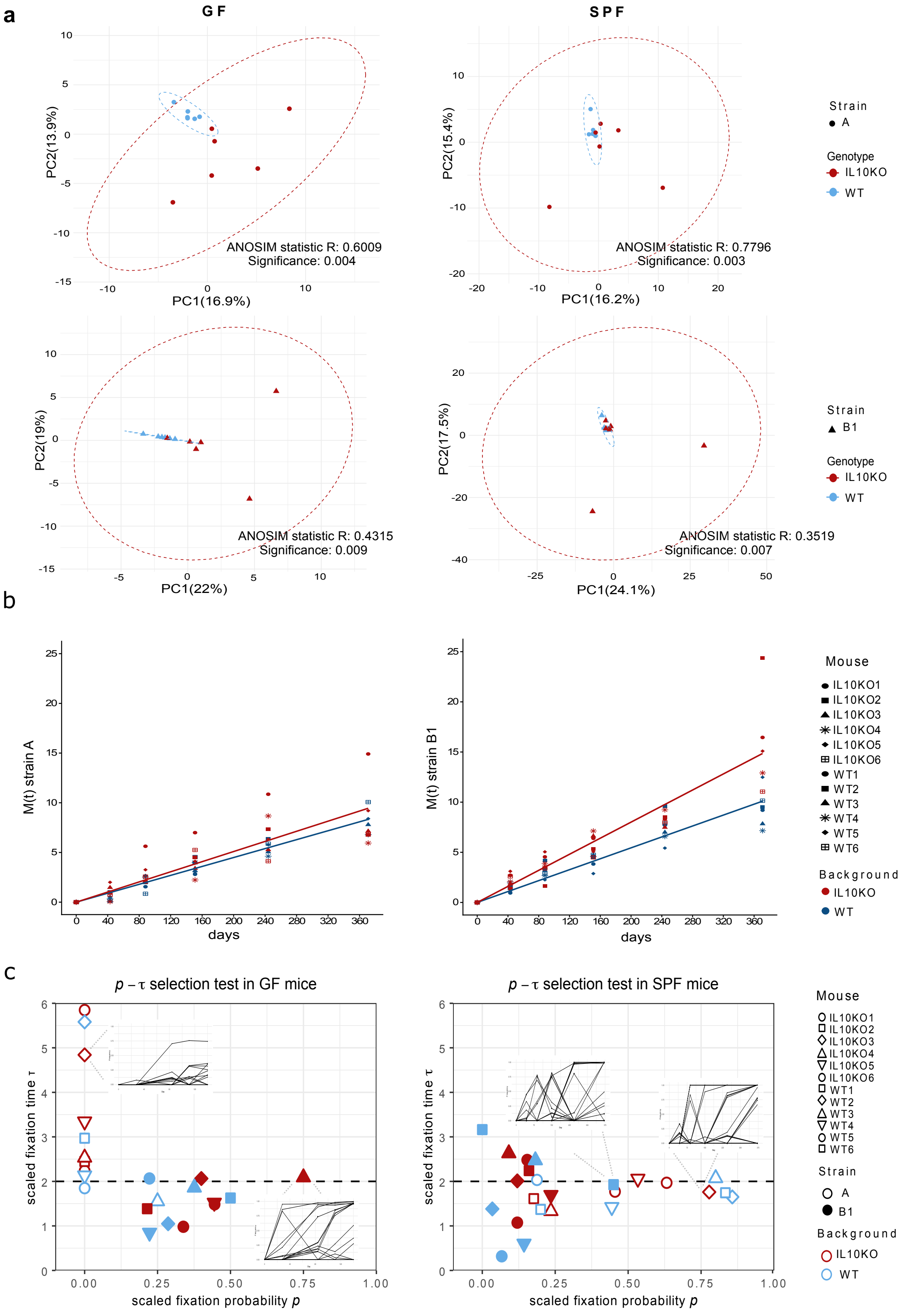


**Supplementary Figure 2. Mutational landscape is dependent of the strain and the mouse background a,** PCA analysis of mutations in each strain depending on host (WT *vs* IL10KO), shows a significant separation of the mutational landscape of each strain with the mouse genetic background. **b,** Mutation accumulation in SPF mice until day 371. Data points represent measured values of M(t), the sum of allele frequencies at each timepoint. Linear models were fitted to strain A and strain B1 data, respectively. **c,** $p$-$\tau$ plot on the frequency and time distributions of mutations in GF and SPF mice, with examples of dynamics of mutations frequencies along time of strain A in GF mice IL10KOM3 and strainB1 in IL10KOM4, and in SPF mice strain A in IL10KOM3 and strain B1 in WT6. $p$-$\tau$ selection test captures the joint statistics of fixation probabilities and time. Data shows that strain A evolution is dominated by diversifying selection in both GF IL10KO and WT mice while strain B1 evolution is dominated by directional selection and clonal interference during evolution in GF IL10KO and WT mice. In the presence of microbiota (SPF) strain A mutational dynamics is consistent with directional selection in both IL10KO and WT mice while for strain B1 both directional selection and diversifying selection shape its evolution.

### Mutational dynamics of strain A in GF mice

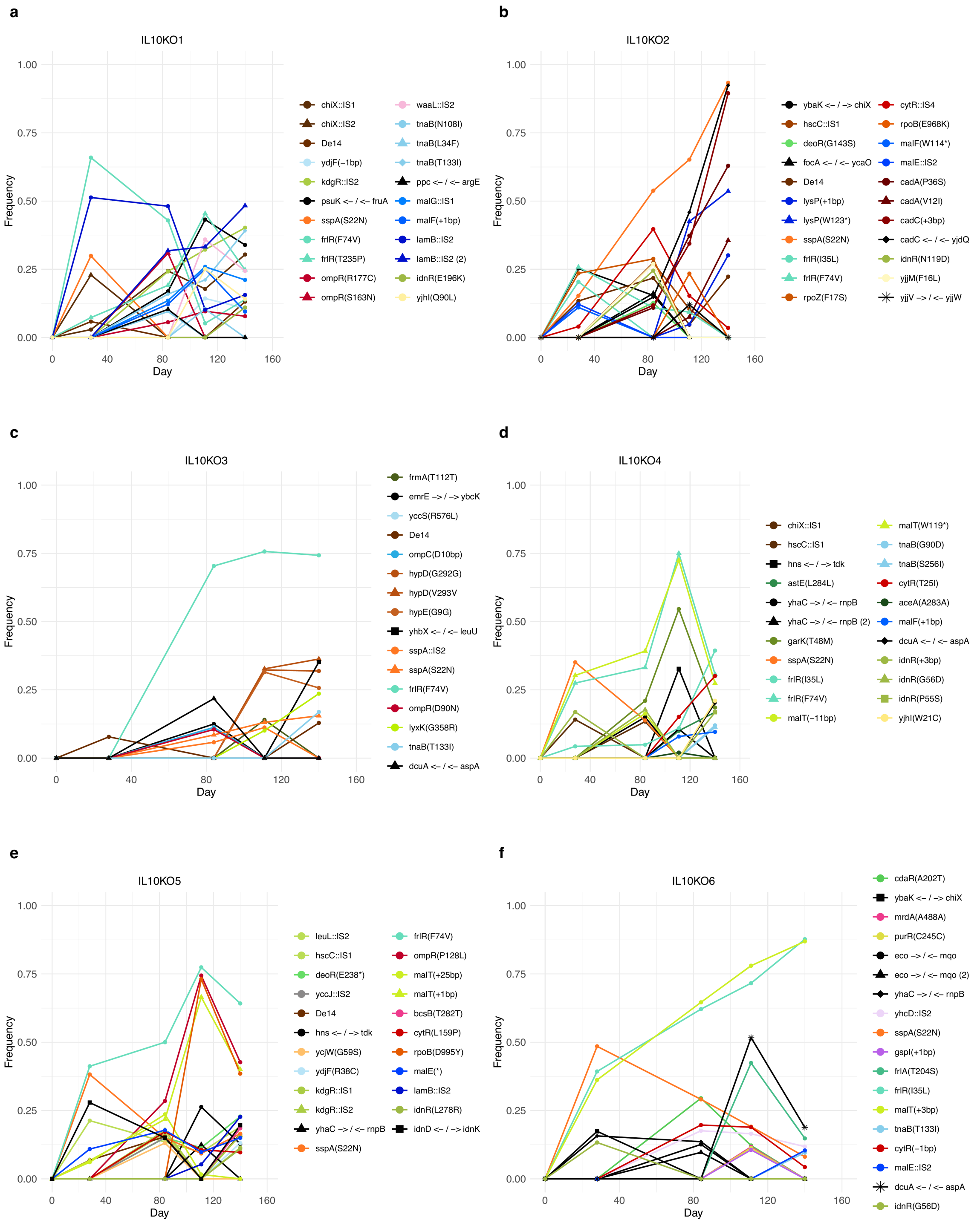

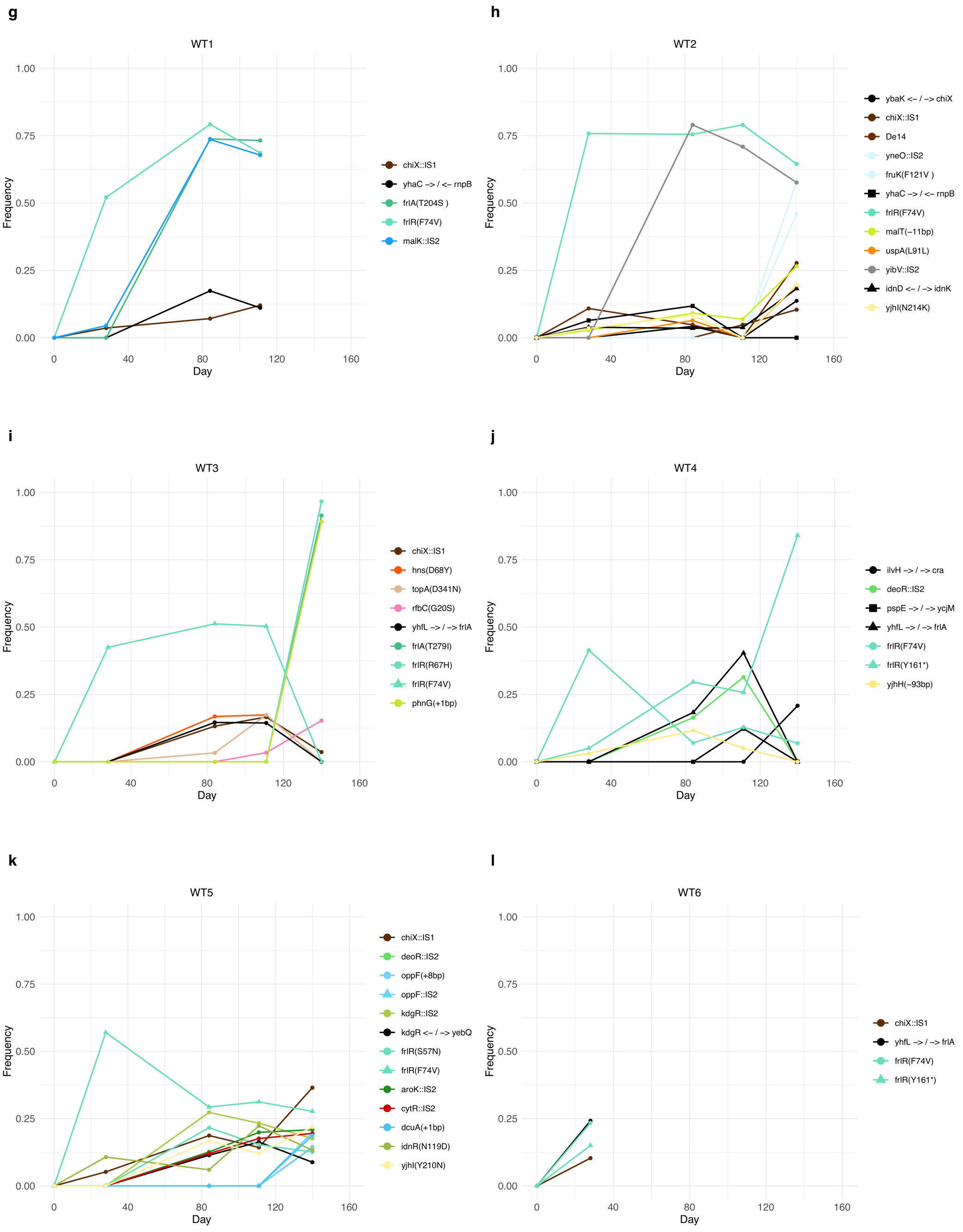


**Supplementary Figure 3. Mutational dynamics of strain A in GF mice.** Frequency dynamics of the mutations identified in strain A across time in GF mouse **a,** IL10KO1, **b,** IL10KO2, **c,** IL10KO3, **d,** IL10KO4, **e,** IL10KO5, **f,** IL10KO6, **g,** WT1, **h,** WT2, **i,** WT3, **j,** WT4, **k,** WT5, **l,** WT6. Legend coloured by function: red/orange – stress response, green/yellow - metabolism, blue - transporters, black - intergenic regions, pink - membrane, purple - adhesion, brown - others, unknow - grey.

#### Mutational dynamics of strain A in SPF mice

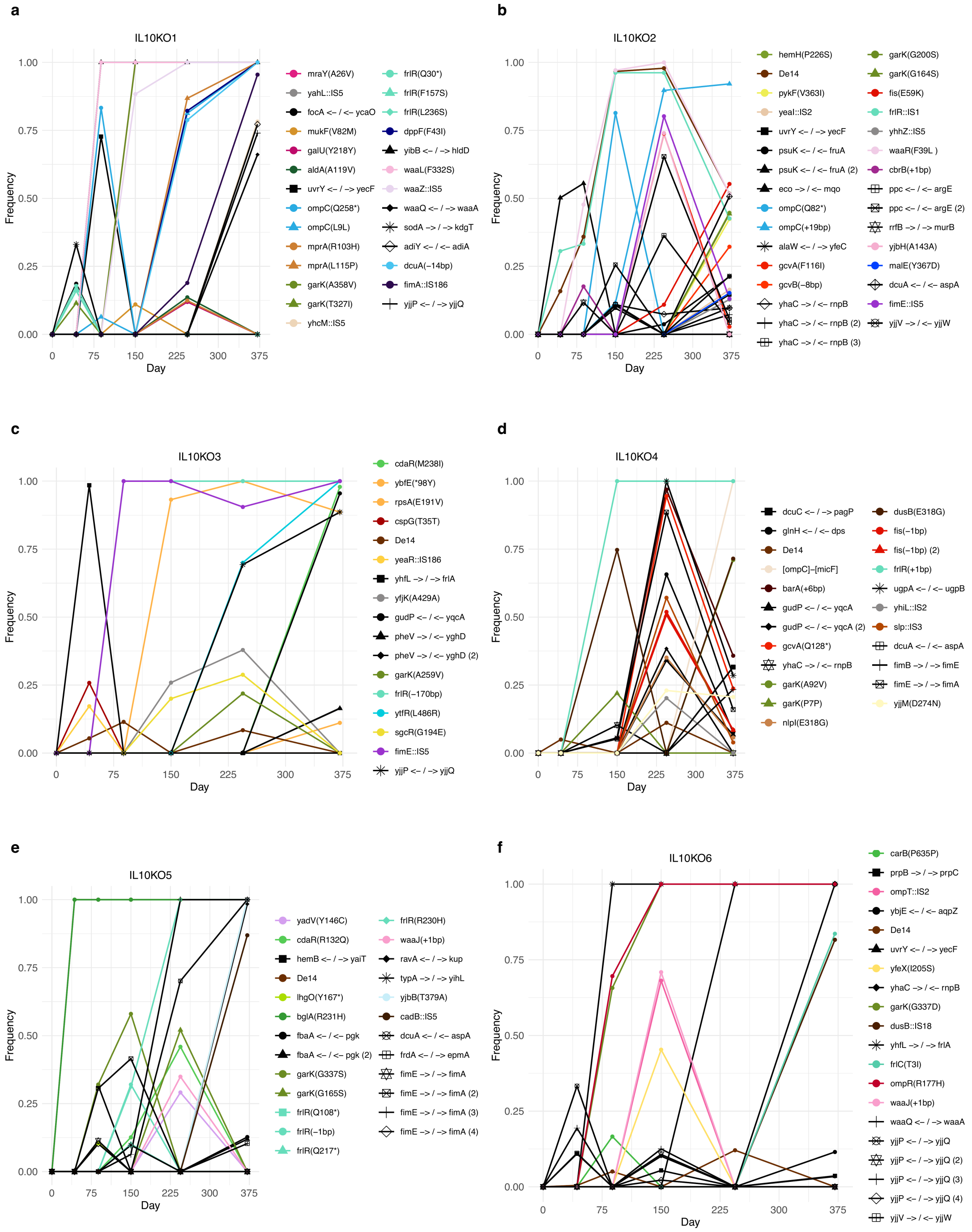

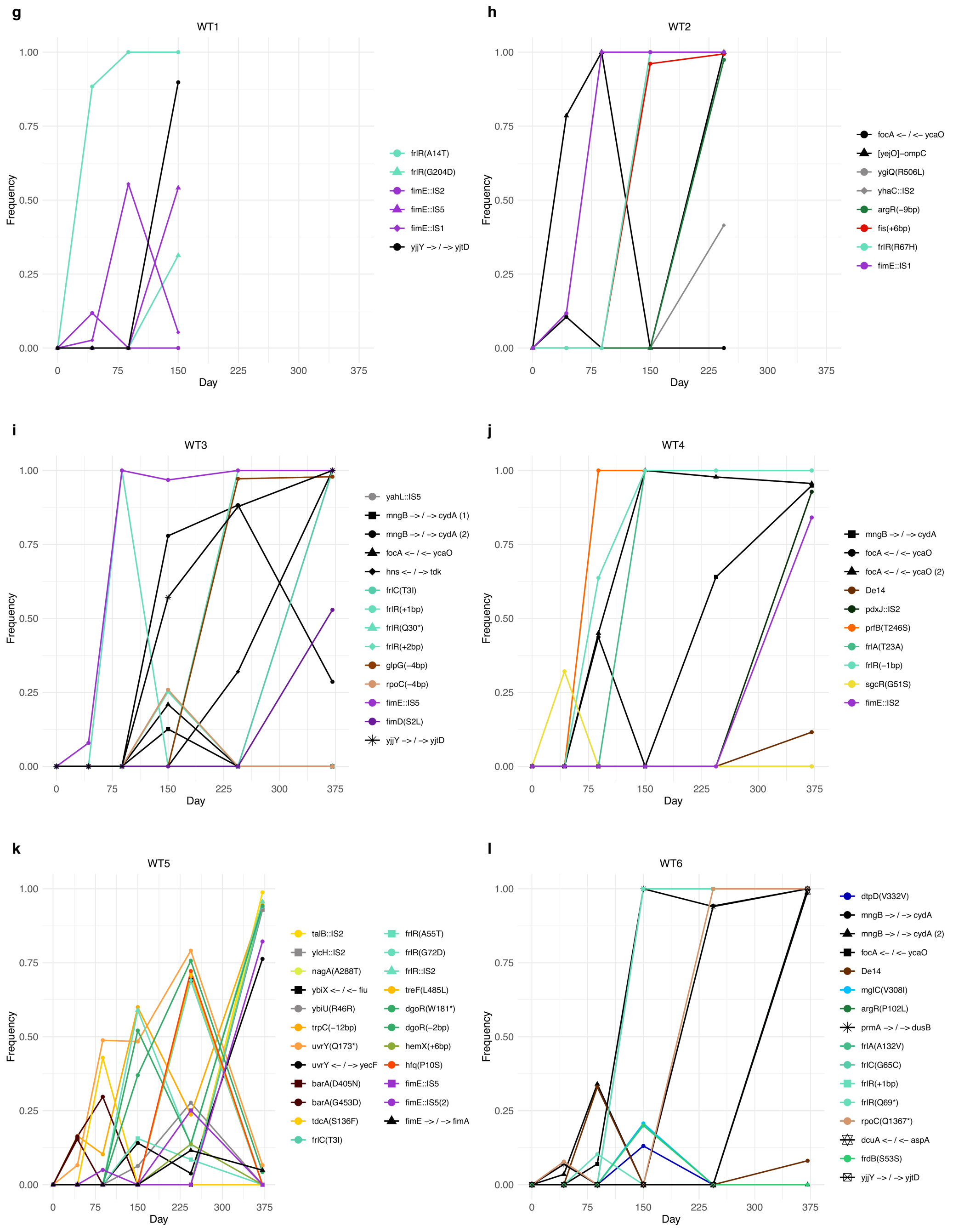


**Supplementary Figure 4. Mutational dynamics of strain A in SPF mice.** Frequency dynamics of the mutations identified in strain A population across time in SPF mouse **a,**IL10KO1, **b,**IL10KO2, **c,**IL10KO3, **d,**IL10KO4, **e,**IL10KO5, **f,**IL10KO6, **g,**WT1, **h,**WT2, **i,**WT3, **j,**WT4, **k,**WT5, **l,**WT6. Legend coloured by function: red/orange – stress response, green/yellow - metabolism, blue - transporters, black - intergenic regions, pink - membrane, purple - adhesion, brown - others, unknow - grey.

#### Mutational dynamics of strain B1 in GF mice

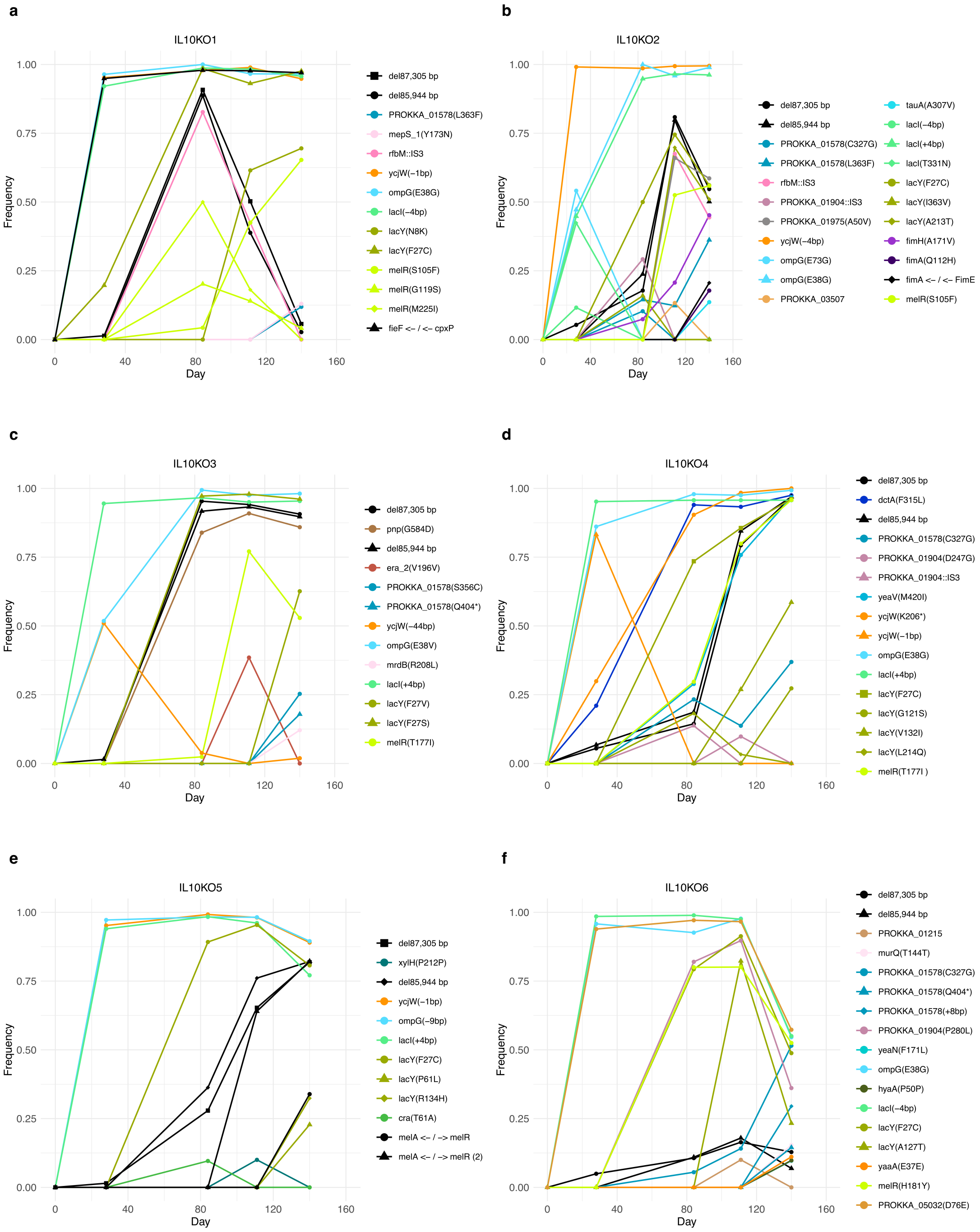

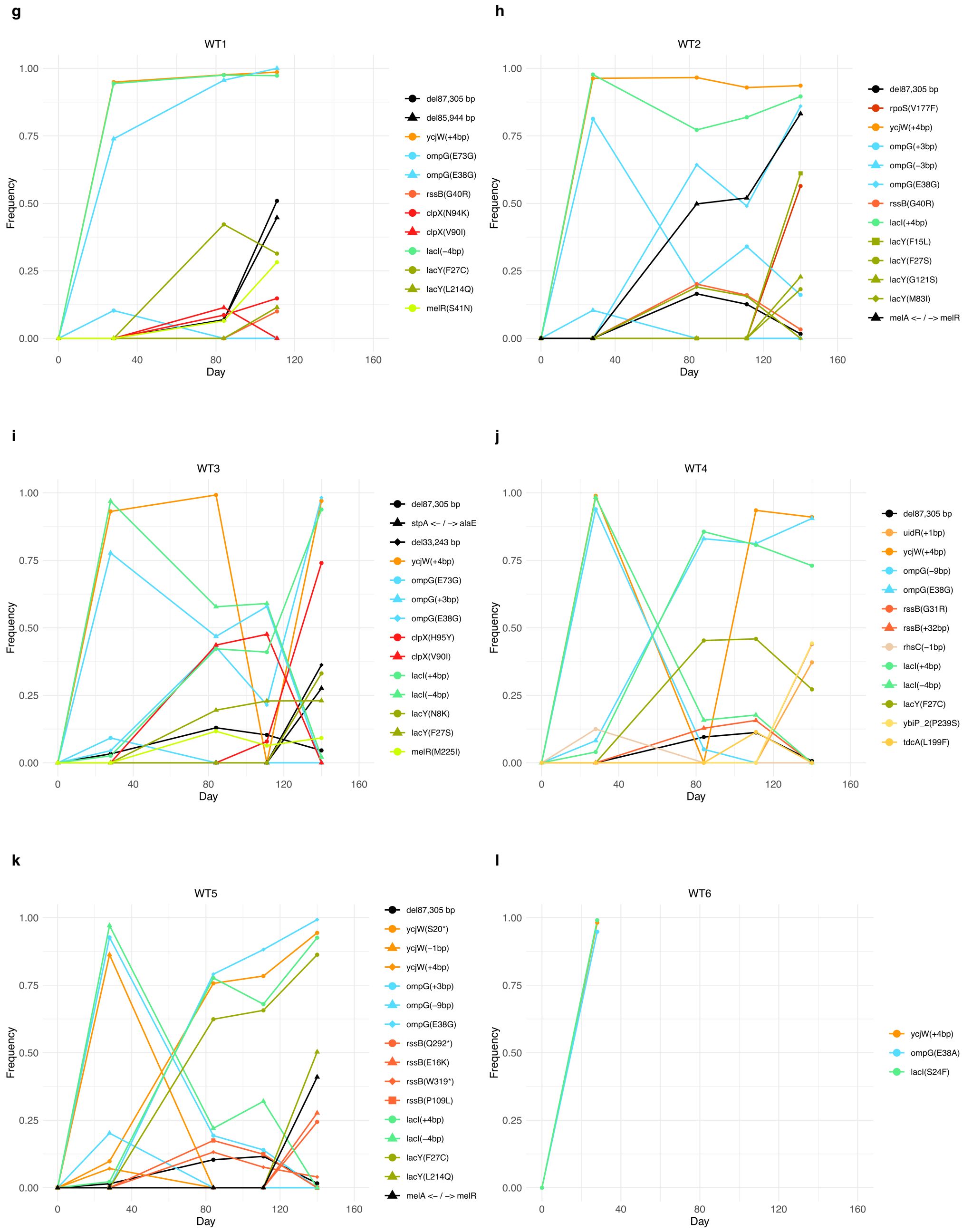


**Supplementary Figure 5. Mutational dynamics of strain B1 in GF mice.** Frequency dynamics of the mutations identified in strain B1 population across time in GF mouse **a,**IL10KO1, **b,**IL10KO2, **c,**IL10KO3, **d,**IL10KO4, **e,**IL10KO5, **f,**IL10KO6, **g,**WT1, **h,**WT2, **i,**WT3, **j,**WT4, **k,**WT5, **l,**WT6. Legend coloured by function: red/orange – stress response, green/yellow - metabolism, blue - transporters, black - intergenic regions, pink - membrane, purple - adhesion, brown - others, unknow - grey.

### Mutational dynamics of strain B1 in SPF mice

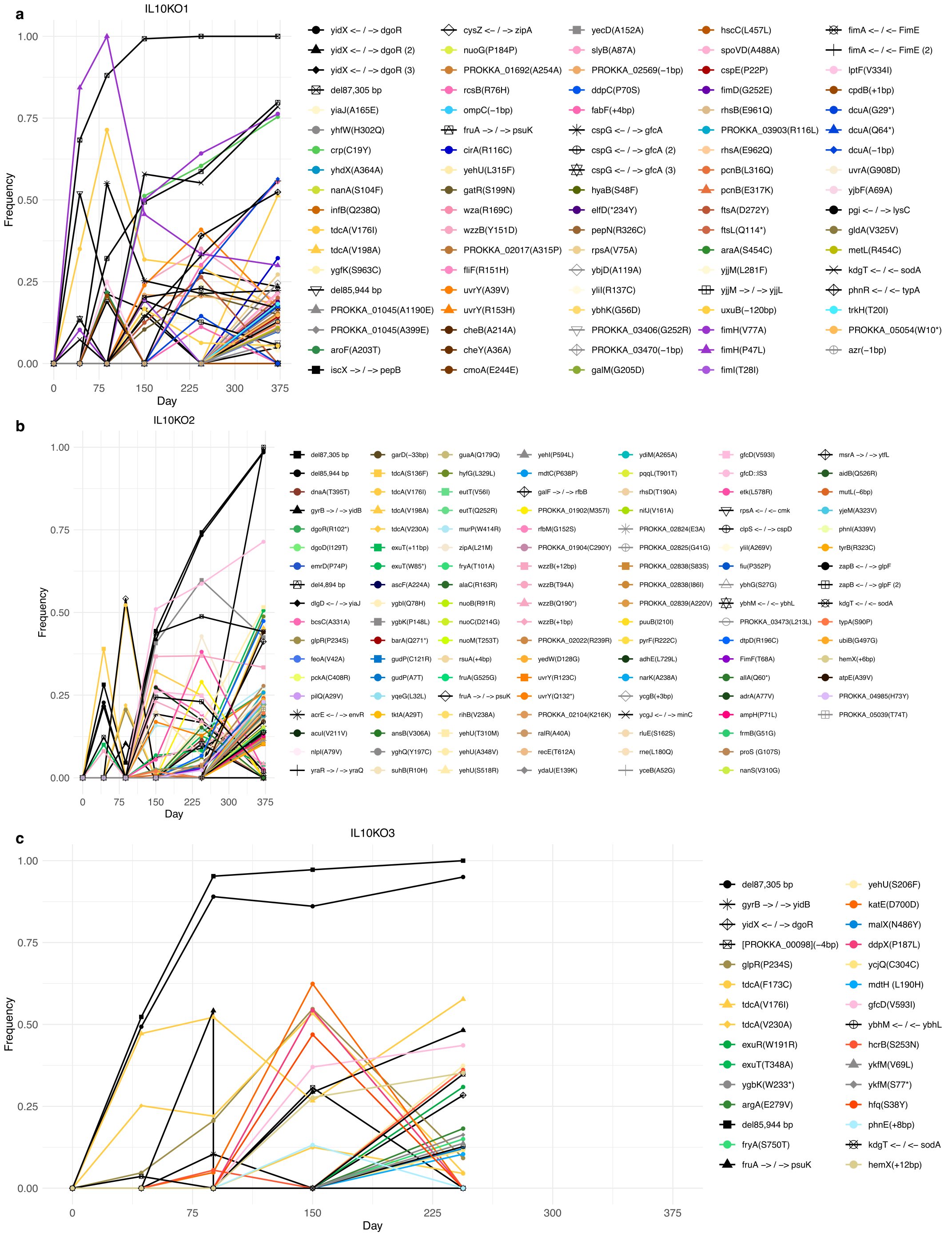

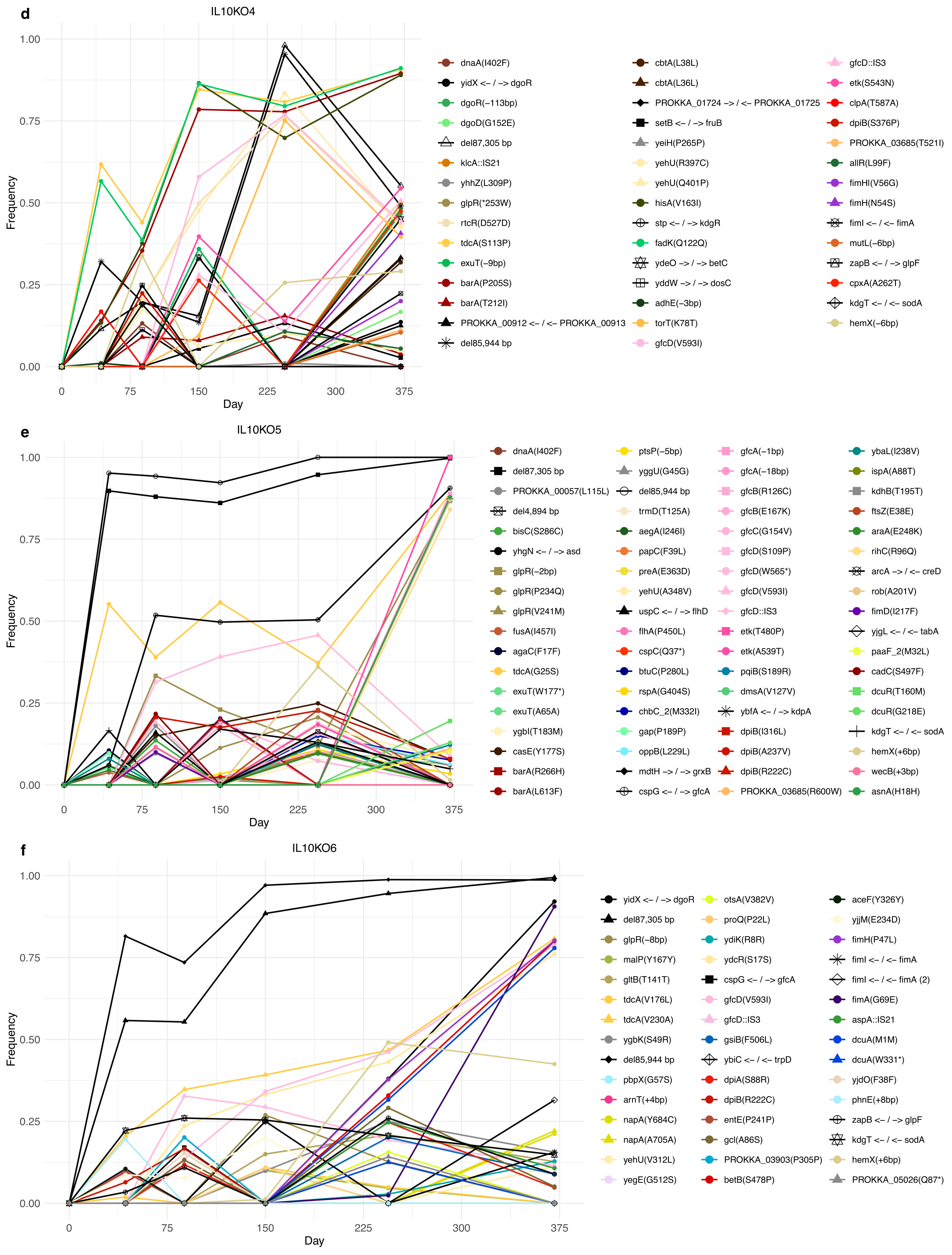
d
IL10KO4
Frequency
Day
1.00
0.75
0.50
0.25
0.00
0
75
150
225
300
375
dnaA(I402F)
yidX <– / –> dgoR
dgoR(−113bp)
dgoD(G152E)
del87,305 bp
klcA::IS21
yhhZ(L309P)
glpR(*253W)
rtcR(D527D)
tdcA(S113P)
exuT(−9bp)
barA(P205S)
barA(T212I)
PROKKA_00912 <– / <– PROKKA_00913
del85,944 bp
cbtA(L38L)
cbtA(L36L)
PROKKA_01724 –> / <– PROKKA_01725
setB <– / –> fruB
yeiH(P265P)
yehU(R397C)
yehU(Q401P)
hisA(V163I)
stp <– / –> kdgR
fadK(Q122Q)
ydeO –> / –> betC
yddW –> / –> dosC
adhE(−3bp)
torT(K78T)
gfcD(V593I)
gfcD::IS3
etk(S543N)
clpA(T587A)
dpiB(S376P)
PROKKA_03685(T521I)
allR(L99F)
fimHI(V56G)
fimH(N54S)
fimI <– / <– fimA
mutL(−6bp)
zapB <– / –> glpF
cpxA(A262T)
kdgT <– / <– sodA
hemX(−6bp)
e
IL10KO5
Frequency
Day
1.00
0.75
0.50
0.25
0.00
0
75
150
225
300
375
dnaA(I402F)
del87,305 bp
PROKKA_00057(L115L)
del4,894 bp
bisC(S286C)
yhgN <– / –> asd
glpR(−2bp)
glpR(P234Q)
glpR(V241M)
fusA(I457I)
agaC(F17F)
tdcA(G25S)
exuT(W177*)
exuT(A65A)
ygbI(T183M)
casE(Y177S)
barA(R266H)
barA(L613F)
ptsP(−5bp)
yggU(G45G)
del85,944 bp
trmD(T125A)
aegA(I246I)
papC(F39L)
preA(E363D)
yehU(A348V)
uspC <– / –> flhD
flhA(P450L)
cspC(Q37*)
btuC(P280L)
rspA(G404S)
chbC_2(M332I)
gap(P189P)
oppB(L229L)
mdtH –> / –> grxB
cspG <– / –> gfcA
gfcA(−1bp)
gfcA(−18bp)
gfcB(R126C)
gfcB(E167K)
gfcC(G154V)
gfcD(S109P)
gfcD(W565*)
gfcD(V593I)
gfcD::IS3
etk(T480P)
etk(A539T)
pqiB(S189R)
dmsA(V127V)
ybfA <– / –> kdpA
dpiB(I316L)
dpiB(A237V)
dpiB(R222C)
PROKKA_03685(R600W)
ybaL(I238V)
ispA(A88T)
kdhB(T195T)
ftsZ(E38E)
araA(E248K)
rihC(R96Q)
arcA –> / <– creD
rob(A201V)
fimD(I217F)
yjgL <– / <– tabA
paaF_2(M32L)
cadC(S497F)
dcuR(T160M)
dcuR(G218E)
kdgT <– / <– sodA
hemX(+6bp)
wecB(+3bp)
asnA(H18H)
f
IL10KO6
Frequency
Day
1.00
0.75
0.50
0.25
0.00
0
75
150
225
300
375
yidX <– / –> dgoR
del87,305 bp
glpR(−8bp)
malP(Y167Y)
gltB(T141T)
tdcA(V176L)
tdcA(V230A)
ygbK(S49R)
del85,944 bp
pbpX(G57S)
arnT(+4bp)
napA(Y684C)
napA(A705A)
yehU(V312L)
yegE(G512S)
otsA(V382V)
proQ(P22L)
ydiK(R8R)
ydcR(S17S)
cspG <– / –> gfcA
gfcD(V593I)
gfcD::IS3
gsiB(F506L)
ybiC <– / <– trpD
dpiA(S88R)
dpiB(R222C)
entE(P241P)
gcl(A86S)
PROKKA_03903(P305P)
betB(S478P)
aceF(Y326Y)
yjjM(E234D)
fimH(P47L)
fimI <– / <– fimA
fimI <– / <– fimA (2)
fimA(G69E)
aspA::IS21
dcuA(M1M)
dcuA(W331*)
yjdO(F38F)
phnE(+8bp)
zapB <– / –> glpF
kdgT <– / <– sodA
hemX(+6bp)
PROKKA_05026(Q87*)

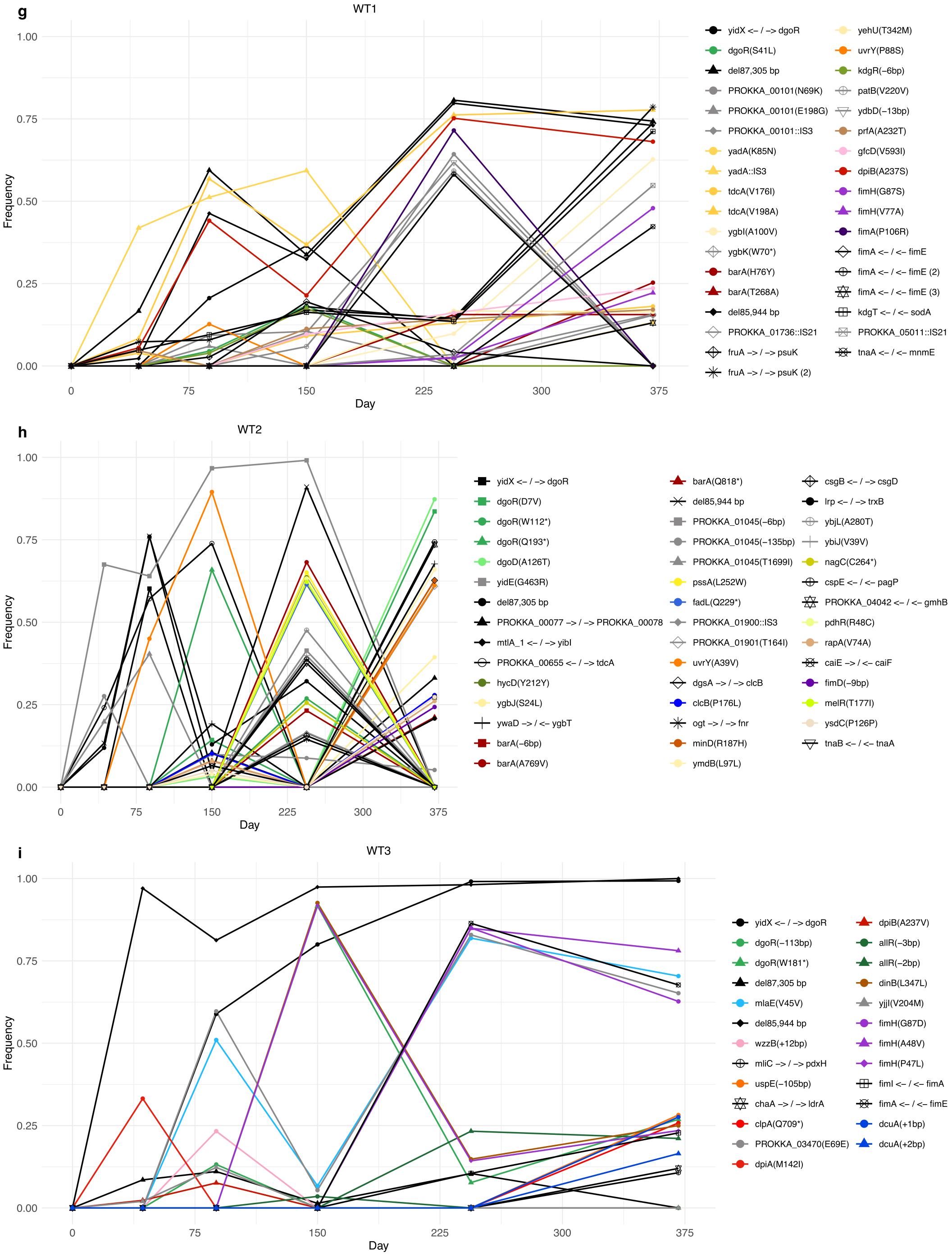

g
WT1
Frequency
Day
yidX <– / –> dgoR
dgoR(S41L)
del87,305 bp
PROKKA_00101(N69K)
PROKKA_00101(E198G)
PROKKA_00101::IS3
yadA(K85N)
yadA::IS3
tdcA(V176I)
tdcA(V198A)
ygbI(A100V)
ygbK(W70*)
barA(H76Y)
barA(T268A)
del85,944 bp
PROKKA_01736::IS21
fruA –> / –> psuK
fruA –> / –> psuK (2)
yehU(T342M)
uvrY(P88S)
kdgR(–6bp)
patB(V220V)
ydbD(–13bp)
prfA(A232T)
gfcD(V593I)
dpiB(A237S)
fimH(G87S)
fimH(V77A)
fimA(P106R)
fimA <– / <– fimE
fimA <– / <– fimE (2)
fimA <– / <– fimE (3)
kdgT <– / <– sodA
PROKKA_05011::IS21
tnaA <– / <– mnmE
h
WT2
Frequency
Day
yidX <– / –> dgoR
dgoR(D7V)
dgoR(W112*)
dgoR(Q193*)
dgoD(A126T)
yidE(G463R)
del87,305 bp
PROKKA_00077 –> / –> PROKKA_00078
mtlA_1 <– / –> yibI
PROKKA_00655 <– / –> tdcA
hycD(Y212Y)
ygbJ(S24L)
ywaD –> / <– ygbT
barA(–6bp)
barA(A769V)
barA(Q818*)
del85,944 bp
PROKKA_01045(–6bp)
PROKKA_01045(–135bp)
PROKKA_01045(T1699I)
pssA(L252W)
fadL(Q229*)
PROKKA_01900::IS3
PROKKA_01901(T164I)
uvrY(A39V)
dgsA –> / –> clcB
clcB(P176L)
ogt –> / –> fnr
minD(R187H)
ymdB(L97L)
csgB <– / –> csgD
lrp <– / –> trxB
ybjL(A280T)
ybiJ(V39V)
nagC(C264*)
cspE <– / <– pagP
PROKKA_04042 <– / <– gmhB
pdhR(R48C)
rapA(V74A)
caiE –> / <– caiF
fimD(–9bp)
melR(T177I)
ysdC(P126P)
tnaB <– / <– tnaA
i
WT3
Frequency
Day
yidX <– / –> dgoR
dgoR(–113bp)
dgoR(W181*)
del87,305 bp
mlaE(V45V)
del85,944 bp
wzzB(+12bp)
mliC –> / –> pdxH
uspE(–105bp)
chaA –> / –> ldrA
clpA(Q709*)
PROKKA_03470(E69E)
dpiA(M142I)
dpiB(A237V)
allR(–3bp)
allR(–2bp)
dinB(L347L)
yjjI(V204M)
fimH(G87D)
fimH(A48V)
fimH(P47L)
fimI <– / <– fimA
fimA <– / <– fimE
dcuA(+1bp)
dcuA(+2bp)

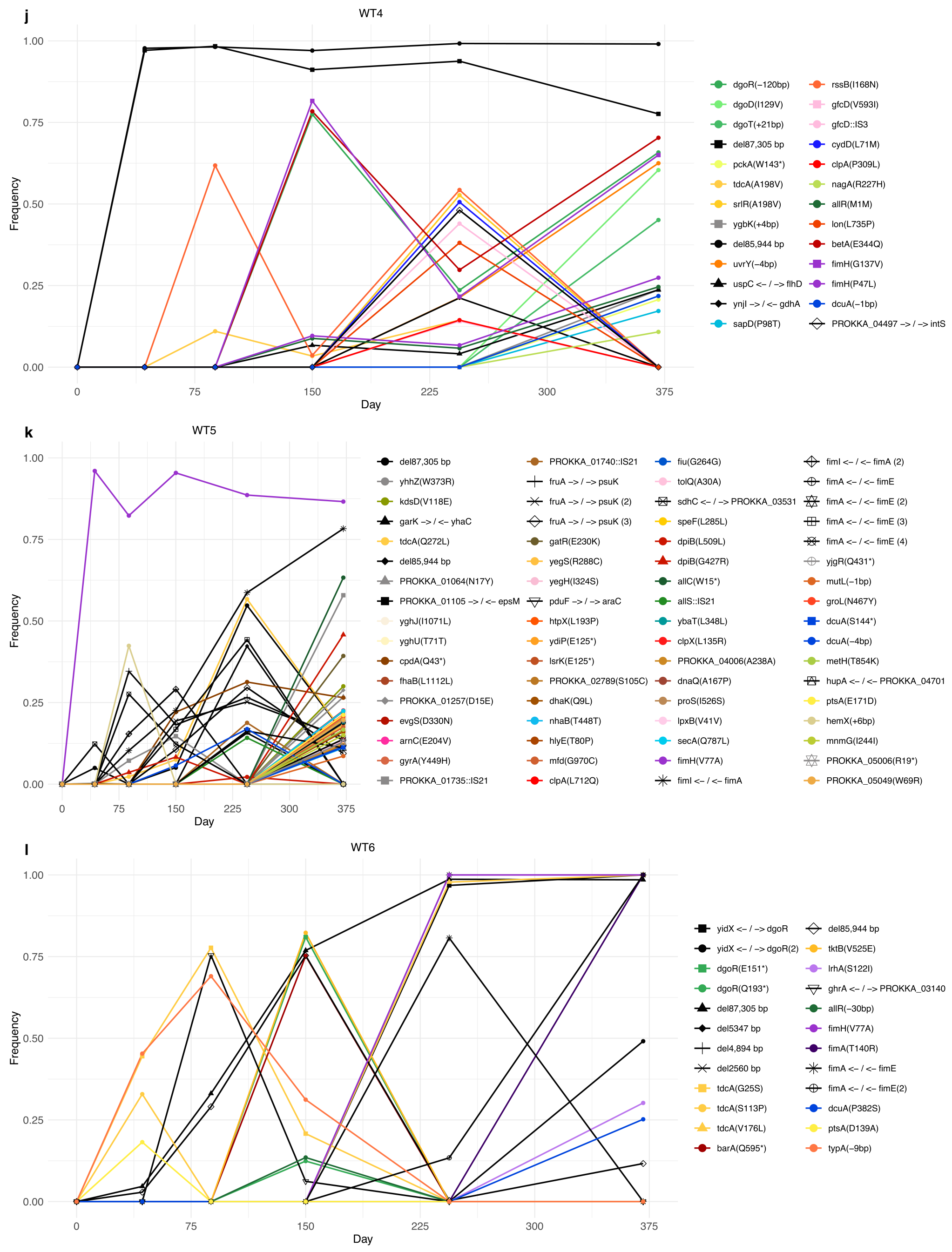


**Supplementary Figure 6. Mutational dynamics of strain B1 in SPF mice.** Frequency dynamics of the mutations identified in strain B1 population across time in SPF mouse **a,**IL10KO1, **b,**IL10KO2, **c,**IL10KO3, **d,**IL10KO4, **e,**IL10KO5, **f,**IL10KO6, **g,**WT1, **h,**WT2, **i,**WT3, **j,**WT4, **k,**WT5, **l,**WT6. Legend coloured by function: red/orange – stress response, green/yellow - metabolism, blue - transporters, black - intergenic regions, pink - membrane, purple - adhesion, brown - others, unknow - grey.

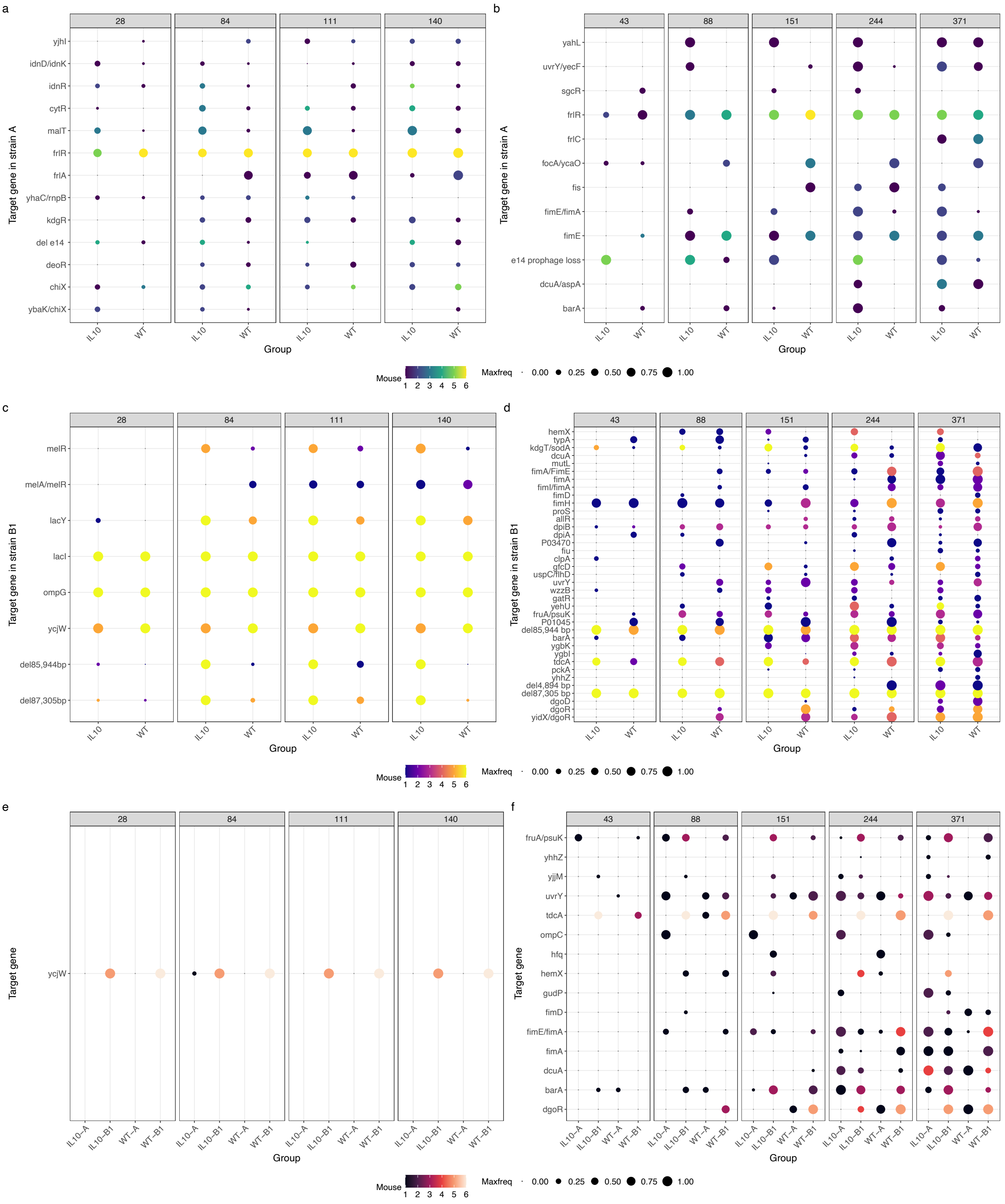


**Supplementary Figure 7. Adaptive targets emerging independently of the inflammatory status of the host** in strain A colonizing **a,** GF and **b,** SPF mice, and in strain B1 colonizing **c,** GF and **d,** SPF mice. Convergent adaptive targets emerging in both strain A and B1 in **e,** GF and **f,** SPF mice. Colours represent the number of mice that acquired mutation in that target gene until that specific timepoint, and circle size represent the maximum frequency reached by that mutation on that timepoint.

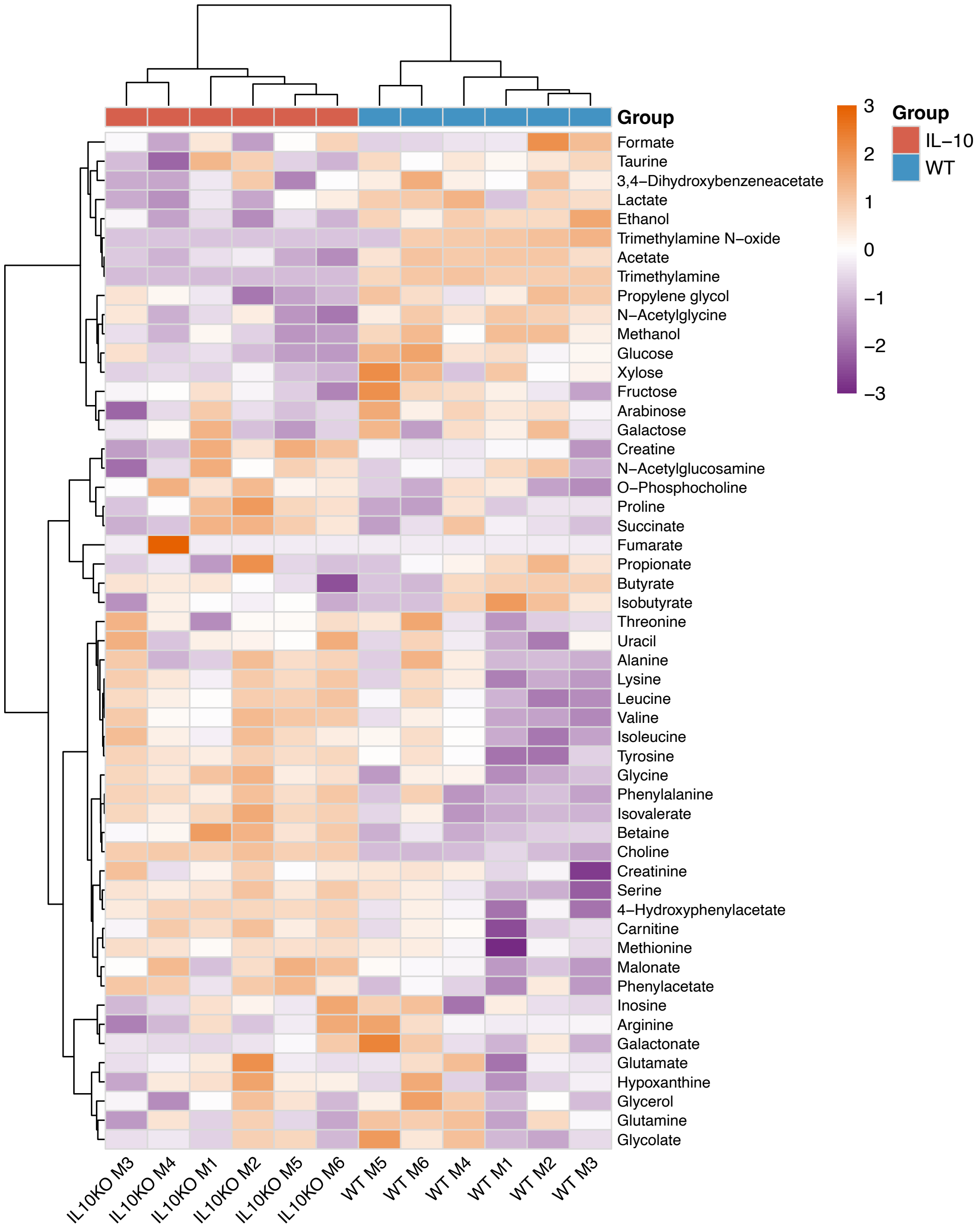


**Supplementary Figure 8. Heatmap of median concentrations of metabolites across the IL10KO and WT SPF mice.** Colours represent indicate row-scaled metabolite abundances (Z-scores). following hierarchical clustering was performed using correlation distance with Ward’s linkage (Ward.D2). The absolute concentration of the different metabolites for each individual mouse is detailed in Supplementary Table 13.

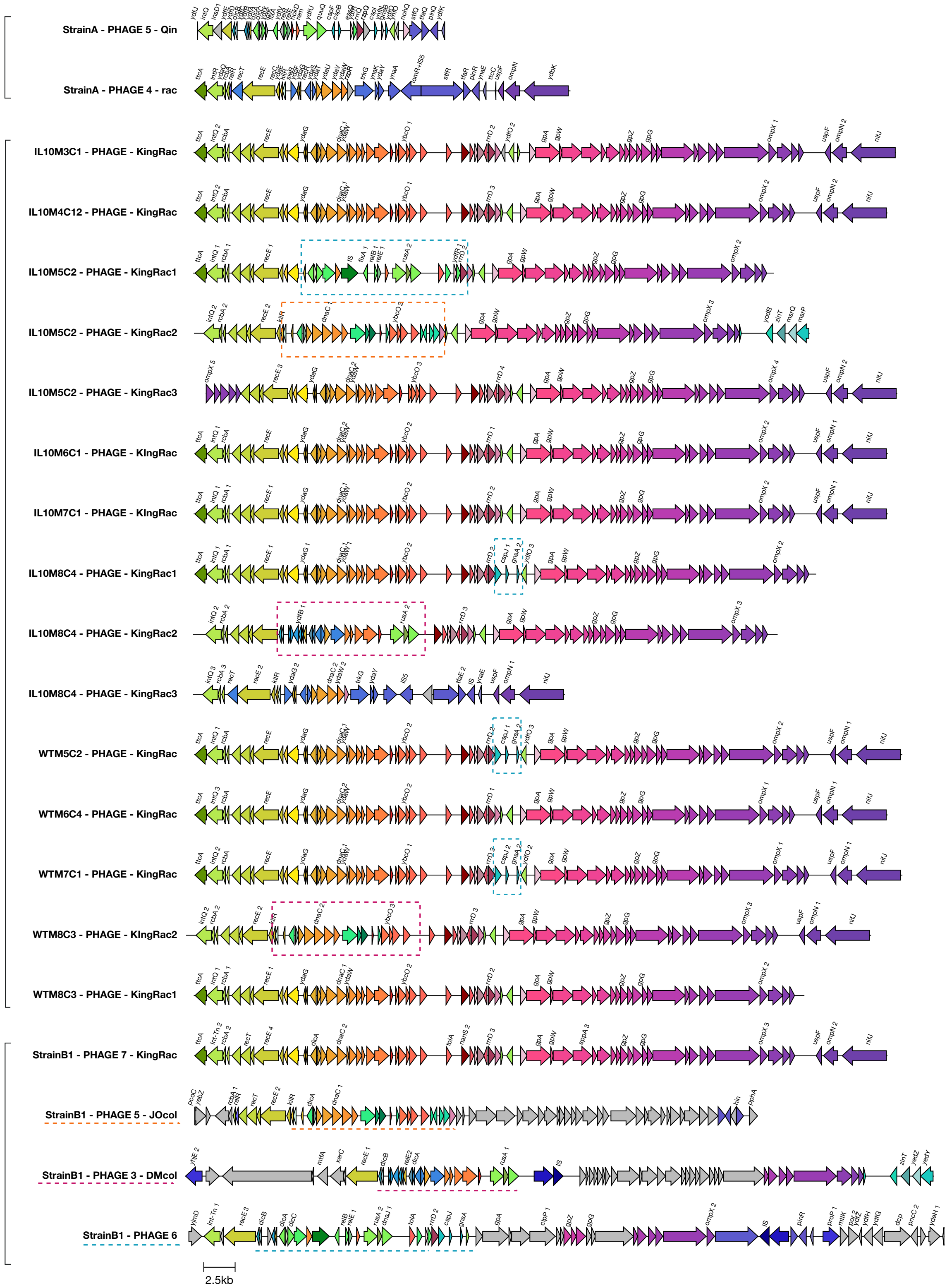


**Supplementary Figure 9. Prophage mosaicism.** KingRac region of several strain A evolved clones, isolated at day 371 of colonization. Some of the isolated clones carry more than one copy of KingRac prophage. Top two prophages are cryptic prophages rac and Qin of ancestral strain A. Bottom four prophages are prophage KingRac, JOcol, DMcol and prophage 6 from strain B1. Orange box represents recombinations of KingRac with Jocol prophage, pink box represents recombination regions with DMcol prophage and blue box represents recombination regions with prophage 6. Visualization was performed using clinker (https://github.com/gamcil/clinker).

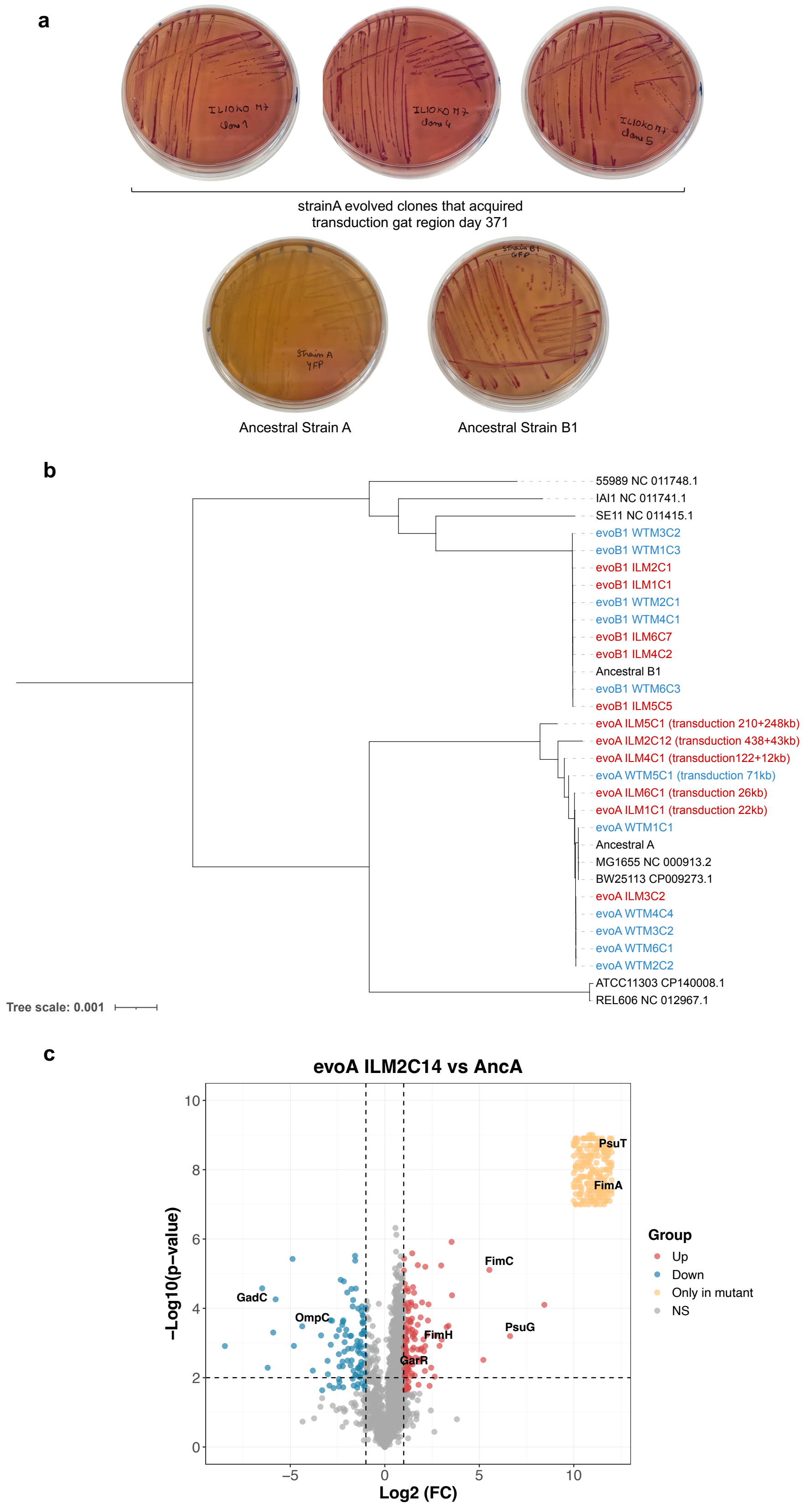


**Supplementary Figure 10. HGT leads to *E. coli* diversification under chronic inflammation. a,** Phenotypic assay showing strain A evolved clones acquiring the ability to consume galactitol, by plating in MacConkey plates supplemented with galactitol. While strain A ancestor does not exhibited changes in colony colour due to absence of galactitol metabolism, its evolved clones from IL10KO mouse 5 show red colonies, as well as the strain B1 ancestor. **b,** Phylogenetic tree including evolved clones from strain A and strain B1 at day 371 (except WT mouse 1 and 2, days 151 and 244 respectively). Phylogenetic analysis performed taking strain A ancestral genome as reference. Results show increase in divergence of clones evolved in SPF IL10KO mice depending on the size of the transduction regions acquired. Clones evolved in IL10KO SPF mice are in red and WT SPF mice are in blue, in black reference/natural isolates from *E. coli* phylogroups A and B1. **c**, Hybrid clone evolved in IL10KO (evoA ILM2C14) shows a significantly changed proteome: proteins of type I fimbriae have higher concentration in the evolved clone compared with its ancestor, as well as PsuG and PsuT involved in pseudouridine degradation and GarR semialdehyde reductase induced by D-glucarate, D-galactarate, and D-glycerate; one of the major non-specific OM porins OmpC is at lower concentration in the evolved clone. Highlighted in red are proteins with higher concentration in evolved IL10KO mouse 2 clone 14, in blue the proteins with lower concentration and in orange the proteins only detected in the evolved clone (Supplementary table 25).
